# Heat shock response pathways regulate stimulus-specificity and sensitivity of NF-κB signalling to temperature stress

**DOI:** 10.1101/782516

**Authors:** Anna Paszek, Małgorzata Kardyńska, James Bagnall, Jarosław Śmieja, David G. Spiller, Piotr Widłak, Marek Kimmel, Wieslawa Widlak, Pawel Paszek

## Abstract

Ability to adapt to temperature changes trough the Heat Shock Response (HSR) pathways is one of the most fundamental and clinically relevant cellular response systems. Here we report that Heat Shock (HS) induces a temporally-coordinated and stimulus-specific adaptation of the signalling and gene expression responses of the Nuclear Factor κB (NF-κB) transcription factor. We show that exposure of MCF7 breast adenocarcinoma cells to 43°C 1h HS inhibits the immediate signalling response to pro-inflammatory Interleukin 1β (IL1β) and Tumour Necrosis Factor α (TNFα) cytokines. Within 4h after HS treatment IL1β-induced responses return to normal levels, but the recovery of the TNFα-induced responses is delayed. Using siRNA knock-down of Heat Shock Factor 1 and mathematical modelling we show that the stimulus-specificity is conferred via the Inhibitory κB kinase signalosome, with HSR differentially controlling individual cytokine transduction pathways. Finally, using a novel mathematical model we predict and experimentally validate that the HSR cross-talk confers differential cytokine sensitivity of the NF-κB system to a range of physiological and clinically-relevant temperatures. This quantitative understanding of NF-κB and HSR cross-talk mechanisms is fundamentally important for the potential improvement of current hyperthermia protocols.

## Introduction

The evolutionary conserved Heat Shock Response (HSR) system regulates how cells adapt to stress (Morimoto, 1998). HSR involves a large family of molecular chaperons called Heat Shock Proteins (HSP) produced as internal repair mechanisms against thermal damage (Richter et al., 2010). In eukaryotic cells, the HSR response is mediated by the Heat Shock Factor 1 (HSF1) transcription factor (Anckar and Sistonen, 2011). In resting cells, HSF1 monomers are kept in an inactive form via their association with heat shock proteins (Akerfelt et al., 2010). High temperatures cause protein conformational changes that in turn leads to a redistribution of HSPs from complexes with HSF1 towards the damaged proteome to initiate repair (Tang et al., 2015). This results in release of HSF1 monomers and their activation via trimerization and posttranslational modifications (Hentze et al., 2016; Zheng et al., 2016). The activated HSF1 transcription factor then leads to the production of different HSP family members. HSPA1 (HSP70) is thought to be the most robustly activated by HSF1, while others including HSP90, the main inhibitor of HSF1 activity are present at a relatively constant level (Akerfelt et al., 2010). HSPs restore protein homeostasis *via de novo* protein folding and targeted degradation and eventually inhibit HSF1 activity (Labbadia and Morimoto, 2015). They also induce a state of thermotolerance limiting further damage to repeated HS (Kampinga, 1993).

Evolutionary conserved NF-κB regulates the expression of hundreds of genes involved in inflammation as well as control of apoptosis, proliferation, cell adhesion and aging (Perkins, 2012). The NF-κB family includes five proteins, all characterised by the presence of the Rel Homology Domain responsible for dimerization as well as DNA and protein binding, but the ubiquitously expressed p65/p50 heterodimer is thought to be most abundant (Hayden and Ghosh, 2008). The NF-κB system integrates a variety of signals, including proinflammatory cytokines, such as Tumour Necrosis Factor α (TNFα) and Interleukin 1β (IL1β), bacterial products, viruses, foreign DNA/RNA and many others (Perkins, 2007). Selective signal integration relies on activation of the inhibitory κB kinase (IKK), a multiprotein signalosome complex composed of IKKα and IKKβ subunits, required for IκB degradation and NF-κB translocation into the nucleus (Adamson et al., 2016; DeFelice et al., 2019; Shembade et al., 2010; Werner et al., 2005). In resting cells, NF-κB is sequestered in the cytoplasm by association with the Inhibitory κB proteins (IκB), but upon stimulation undergoes nuclear-to-cytoplasmic oscillations as a result of cyclic degradation and NF-κB-dependent resynthesis of IκB and A20 (Hoffmann et al., 2002; Nelson et al., 2004). The timing, e.g. frequency and amplitude of NF-κB was shown to control target gene expression (Ashall et al., 2009; Kellogg and Tay, 2015; Tay et al., 2010), which may confer both proapoptotic or prosurvival functions (Hayden and Ghosh, 2008). NF-κB signalling plays a key role in disease and in particular cancer progression (Perkins, 2004). In breast cancer, the NF-κB activity stimulates tumour growth, metastasis and chemoresistance, therefore therapeutic inhibition of its activity is considered beneficial (Esquivel-Velazquez et al., 2015).

Hyperthermia, the exposure of tissue to high temperature has been considered a promising strategy to sensitise cancer cells to therapeutic intervention (Wust et al., 2002). However, a better understanding of the temperature effect on cellular signalling, in general, and crosstalk between HSR and NF-κB systems, in particular, is required for better efficacy. The NF-κB system can adapt to physiological (<40°C) temperatures (Harper et al., 2018), but exposure to higher temperatures (>40°C) results in the attenuation of the NF-κB signalling and function (Esquivel-Velazquez et al., 2015; Janus et al., 2015; Kardynska et al., 2018; Wong et al., 1997b). This involves IKK denaturation (Kardynska et al., 2018; Lee et al., 2004; Pittet et al., 2005), with inhibition of IκB degradation, NF-κB phosphorylation and translocation as well as target gene expression (Ayad et al., 1998; Cooper et al., 2010; Feinstein et al., 1997; Furuta et al., 2004; Janus et al., 2015; Wong et al., 1997a, b). Mechanistic understanding of underlying processes is confounded by contradictory findings and lack of quantitative analyses. For example, HSPA1 and HSP90 were shown to repair and stabilize IKK (Jiang et al., 2011; Lee et al., 2005), while overexpression of HSPA1 was found to inhibit IKK activity (Sheppard et al., 2014) and TRAF2-mediated NF-κB activation (Dai et al., 2010). Here we use interdisciplinary systems biology approaches to systematically understand how elevated temperature affects NF-κB signalling to cytokine treatment using live single-cell imaging and mathematical modelling.

## Results

### HS modulates the NF-κB signalling responses to TNFα stimulation

Previously we characterised HS-mediated inhibition of the TNFα-dependent NF-κB signalling (Kardynska et al., 2018). Here we sought to investigate the long-term modulation and recovery of NF-κB responses following HS. We used a treatment protocol where breast adenocarcinoma MCF7 cells were exposed to 1h 43°C HS, subsequently recovered in normal conditions (37°C) for up to 4h and treated with 10 ng/ml TNFα (Fig. 1A). This defines a “recovery time” between the end of HS exposure and the cytokine treatment, which is relevant to clinical hyperthermia protocols (Maluta and Kolff, 2015; Rybinski et al., 2013). In cells maintained under normal conditions, we observed a robust Ser536 p65 phosphorylation, a marker of NF-κB activity, which peaked around 15 min after TNFα stimulation (Fig. 1B). In contrast, stimulation immediately after 1h HS exposure resulted in an inhibition of Ser536 p65 phosphorylation. In these cells, the highest level of p65 activation was observed at 60 min after stimulation. Following extended recovery times, Ser536 p65 phosphorylation steadily increased, however, responses did not fully recover even when TNFα was applied 4h after HS exposure. This demonstrates a long-term attenuation of the NF-κB signalling response by HS. HS exposure may lead to heterogenous NF-κB responses in single cells, which can be masked in the population level analyses (Kardynska et al., 2018). Therefore, we developed a MCF7 cell line stably expressing p65-EGFP suitable for quantitative single time-lapse microscopy analyses. We confirmed, that transduced cells exhibited TNFα and HS-mediated Ser536 p65 activation essentially consistent with that of the wild type cells (Fig. S1A), while HS exposure alone did not cause p65-EGFP translocation (Fig. S1B). Using live single-cell microscopy, we assayed TNFα-induced responses after different HS recovery times in the fluorescent reporter line (see Fig. 1C for confocal images of representative cells, 1D and E for analysis of single cell traces). In agreement with previous imaging studies (Sero et al., 2015; Stewart-Ornstein and Lahav, 2016), stimulation of MCF7 cells cultured in normal conditions resulted in a rapid nuclear translocation of p65-EGFP followed by series of damped oscillations. Individual cells exhibited varied amplitude of the first nuclear translocation (mean 101 arbitrary fluorescence units ± 44 standard deviation) as well as degree of damping. Approximately 50% of cells exhibited up to three nuclear NF-κB translocations, while no cells showed more than five translocations within the 10h imaging window (see Fig. 1E for the distribution of number of translocations). TNFα stimulation immediately after exposure to 1h HS resulted in statistically significant inhibition of the p65-EGFP responses where cells exhibited only residual and delayed activity (Fig. 1D, as well as reduced area under the curve, AUC, Fig. 1E). As the recovery time increased, we observed increases in the first peak nuclear p65-EGFP amplitude, which coincided with a faster response time (Fig. 1E). However, even after a 4h recovery period the initial NF-κB response remained suppressed compared to cells cultured in the normal conditions. Interestingly, the HS exposure induced the long-term oscillatory phase of the NF-κB response (as evident by heat-maps and the increased number of cell exhibiting multiple nuclear NF-κB translocations, Fig. 1E). After 4h recovery, almost 90% of cells exhibited at least five nuclear translocations (and 50% after 2h recovery) within 10h after cytokine treatment. This is indicative of HS-mediated response modulation (Harper et al., 2018).

**Figure 1.**
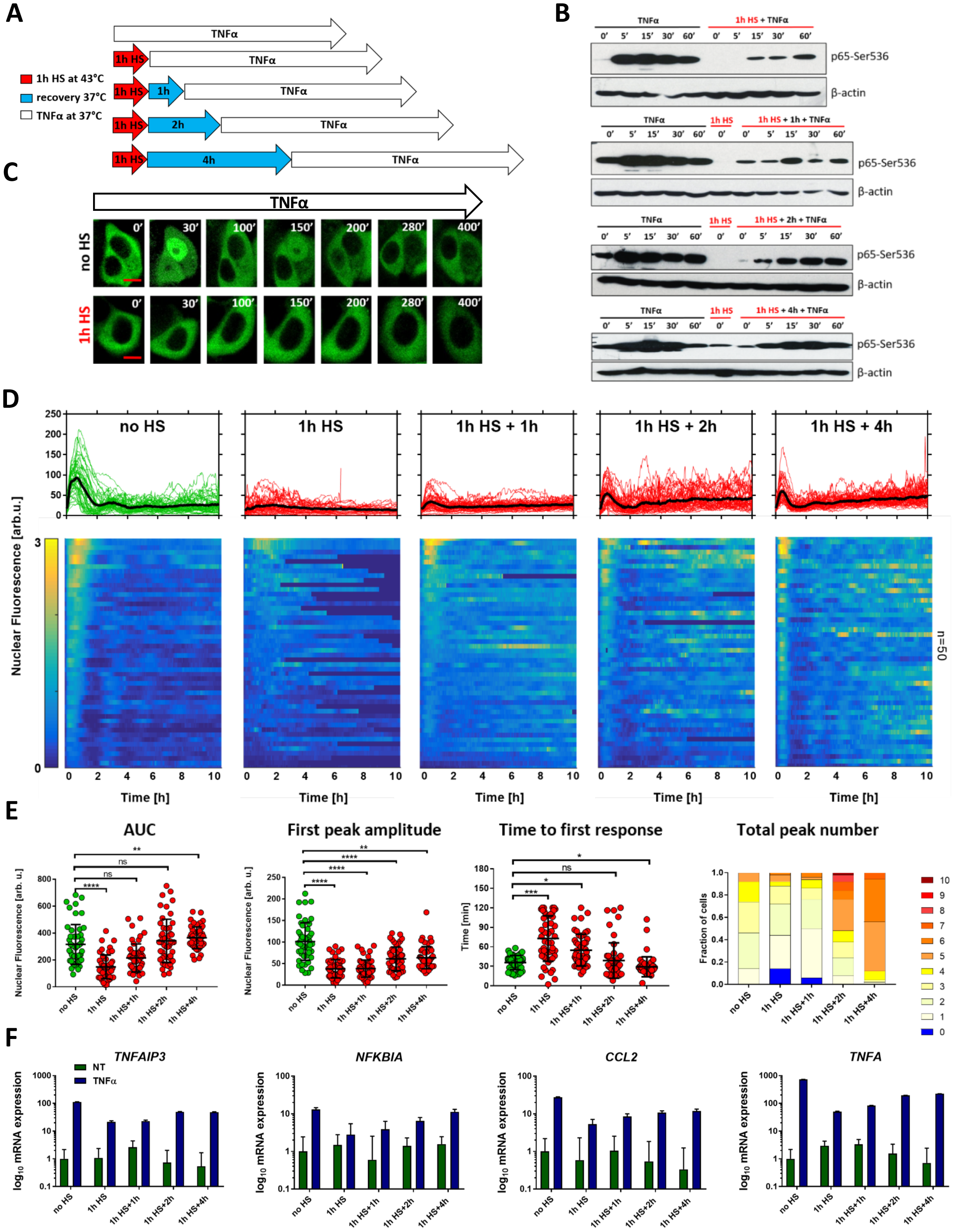
Heat shock modulates NF-κB signalling responses to TNFα stimulation. (**A**) Schematic representation of HS and TNFα treatment: MCF7 cells were either cultured under normal conditions (37°C) or subjected to 1h HS at 43°C, recovered for 0, 1, 2 or 4 hours, and treated with TNFα. (**B**) The level of p65-Ser536 phosphorylation in response to HS (in red) and TNFα treatment (as in A) assayed via Western blotting in whole cell lysates at indicated times (in min). Shown also are untreated controls (0’) and β-actin loading control. (**C**) Confocal microscopy images of representative MCF7 cells stably expressing p65-EGFP. Top panel: cells cultured under normal conditions (at 37°C, no HS) and stimulated with TNFα for indicated time (displayed in minutes). Bottom panel: cells exposed to 1h HS at 43°C prior to TNFα stimulation. Scale bar 5 µm. (**D**) Nuclear NF-κB trajectories in MCF7 cells stably expressing p65-EGFP for different HS and TNFα treatment protocols. Top: individual single cell trajectories (n = 50 per condition, in aribirary fluorescene units) depicted with colour lines; population average – depicted with a black line. Bottom: heat maps of single cell trajectories normalized across all considered conditions (represented on a 0-3 scale). (**E**) Characteristics of single cell NF-κB trajectories from D. From the left: distribution of area under the curve (AUC), first peak amplitude, time to first response and the total peak number. Individual cell data are depicted with circles (with mean ± SD per condition). Kruskal-Wallis one-way ANOVA with Dunn’s multiple comparisons test was used to assess differences between groups (*p<0.05, **p<0.01, ***p<0.001, ****p<0.0001, ns – not significant). (**F**) Quantitative RT-PCR analysis of *TNFAIP3, NFKBIA, CCL2* and *TNFA* mRNA abundance in response to TNFα following HS treatment, in reference to cells cultured at 37°C without TNFα, no HS. Cells assayed at 90 min after TNFα stimulation. Shown are means ± SDs based on three replicate experiments.

In order to understand the functional consequence of the HS-mediated NF-κB inhibition, we measured expression of NF-κB target genes by quantitative RT-PCR at 90 mins after TNFα stimulation (corresponding to the early activation phase, Fig. 1F and Table S1). The analysis of the NF-κB negative inhibitor *TNFAIP3* and *NFKBIA* genes (coding for A20 and IκBα, respectivelly), as well as *TNFA* and chemokine *CCL2* demonstrates a substantial supression of the TNFα-induced mRNA expression immediately after HS exposure. However, following the 4h recovery time, degree of the activation of *TNFAIP3* and *NFKBIA* returned nearly to the levels observed in cells cultured in normal conditions. In contrast, activation of *TNFA* and *CCL2* was still lower (approximately 30%), perhaps reflecting their multifaced regulation (Falvo et al., 2010; Liu et al., 2012).

### Recovery of the HS-mediated NF-κB signalling response is cytokine specific

TNFα acts through its cognate receptor to activate IKK, and a number of mechanisms have been proposed to understand how HSR affect this process (Kardynska et al., 2018; Sheppard et al., 2014). In the first attempt to investigate these mechanisms, we utilised IL1β cytokine, which is known to activate IKK via signal transduction pathway parallel to that of TNFα (Fig. 2A) (Adamson et al., 2016; DeFelice et al., 2019; Shembade et al., 2010; Werner et al., 2008). Using a treatment protocol analogous to that for TNFα (Fig. 2B) we assayed early population-level NF-κB responses to 10 ng/ml IL1β treatment following HS (Fig. 2C). In cells maintained under normal conditions, IL1β treatment induced a rapid induction of Ser536 p65 phosphorylation, which was blocked immediately after HS exposure. However, in contrast to TNFα, IL1β-induced Ser536 phosphorylation levels appeared to return to the pre-HS level after 4h recovery. We performed live single-cell microscopy experiments using the MCF7 cell line stably expressing p65-EGFP fusion protein after treatment with IL1β (Fig. 2D-F). Stimulation of cells maintained in normal conditions resulted in rapid and robust p65-EGFP translocation to the nucleus. In contrast to TNFα treatment, most of the cells (80%) exhibited only a single nuclear translocation, while the remaining 20% of cells exhibited up to four translocations. IL1β treatment immediately after exposure to 1h HS resulted in inhibition of the p65-EGFP response, although some cells exhibited residual but delayed activation (as revealed by a heat map, Fig. 2E). Responses to stimulation after different recovery times revealed a complete transition to pre-HS levels, which was exhibited by increasing first peak amplitude, AUC as well as reducing response time (Fig. 2F). In contrast to TNFα, HS exposure did not alter the long-term NF-κB dynamics, as the distribution of peak numbers did not change compared to cells maintained under normal conditions. Functionally, HS inhibited NF-κB-dependent transcription, which subsequently recovered to the pre-HS levels (albeit to lesser extent for cytokines *TNFA* and *CCL2* than for NF-κB-feedback genes *TNFAIP3* and *NFKBIA*) (Fig. 2G).

**Figure 2.**
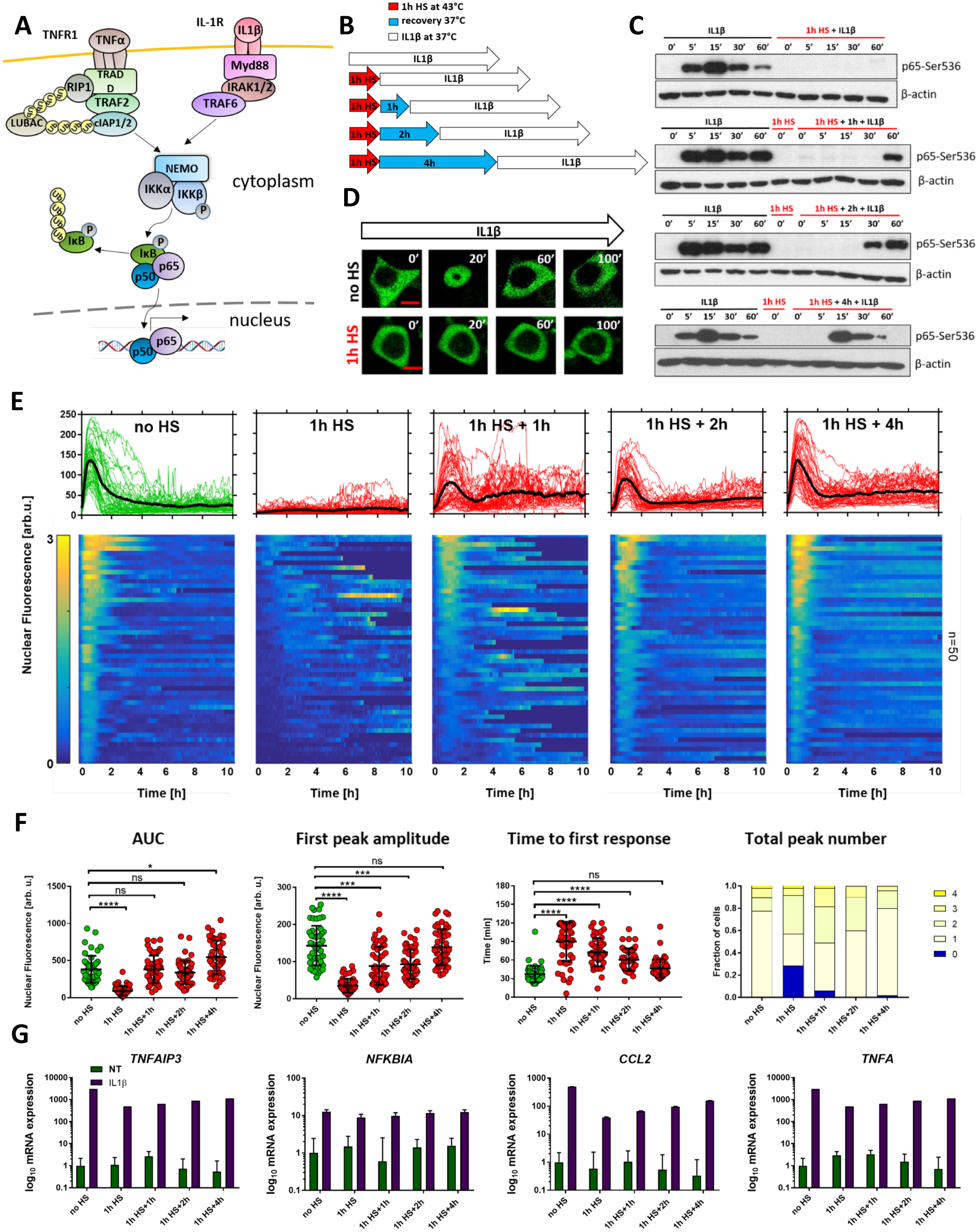
HS-mediated NF-κB signalling response is cytokine dependent. (**A**) Schematic diagram of TNFα and IL1β-dependent signal transduction pathways leading to IKK and NF-κB activation. (**B**) Schematic representation of HS and IL1β treatment: MCF7 cells were either cultured under normal conditions (37°C) or subjected to 1h HS at 43°C, recovered for 0, 1, 2 or 4 hours, and treated with IL1β. (**C**) The level of p65-Ser536 phosphorylation in response to HS (in red) and IL1β (as in A) treatment assayed via Western blotting in whole cell lysates at indicated times (in min). Shown also are untreated controls (0’) and β-actin loading control. (**D**) Confocal microscopy images of representative MCF7 cells stably expressing p65-EGFP. Top panel: cells cultured under normal conditions (at 37°C, no HS) and stimulated with IL1β for indicated time. Bottom panel: cells exposed to 1h HS at 43°C prior to IL1β stimulation. Time after IL1β stimulation displayed in minutes. Scale bar 5 µm. (**E**) Nuclear NF-κB trajectories in MCF7 cells stably expressing p65-EGFP for different HS and IL1β treatment protocols. Top: individual single cell trajectories (n = 50 per condition, in aribirary fluorescene units) depicted with coloured lines; population average – with a black line. Bottom: heat maps of single cell trajectories normalized across all considered conditions (represented on a 0-3 scale). (**F**) Characteristics of single cell NF-κB trajectories from E. From the left: distribution of area under the curve (AUC), first peak amplitude, time to first response and the total peak number. Individual cell data are depicted with circles (with mean ± SD per condition). Kruskal-Wallis one-way ANOVA with Dunn’s multiple comparisons test was used to assess differences between groups (*p<0.05, ***p<0.001, ****p<0.0001, ns – not significant). (**G**) Quantitative RT-PCR analysis of *TNFAIP3, NFKBIA, CCL2* and *TNFA* mRNA abundance in response to HS and IL1β treatment, in reference to cells cultured at 37°C without IL1β, no HS. Cells assayed at 90 min after IL1β stimulation. Shown are means ± SDs based on three replicate experiments.

Overall, these data demonstrate that while the HS exposure may effectively inhibit cytokine-mediated NF-κB signalling and gene expression responses, the kinetic of adaptation to HS is stimulus specific.

### HSF1 differentially regulates cytokine-specific NF-κB signalling

The cellular response to HS involves the HSF1-dependent transcription of genes encoding HSPs as a part of an internal stress-adaptation mechanism (Fig. 3A). Exposure of MCF7 cells to 1h HS resulted in hyperphosphorylation of HSF1 (visible as a shift of the HSF1 protein in western blot analysis which coincided with Ser326 phosphorylation, a marker of temperature-induced HSF1 activation; Fig. 3B) (Guettouche et al., 2005). While transient, the HSF1 activation resulted in a robust upregulation of the HSPA1, a major factor involved in the internal protein repair (Brocchieri et al., 2008). The *HSPA1* mRNA levels showed steady increases for up to several hours after 1h HS exposure (Fig. 3C), while an increased HSPA1 protein level was observed from the second/third hour of recovery (Fig. 3B).

**Figure 3.**
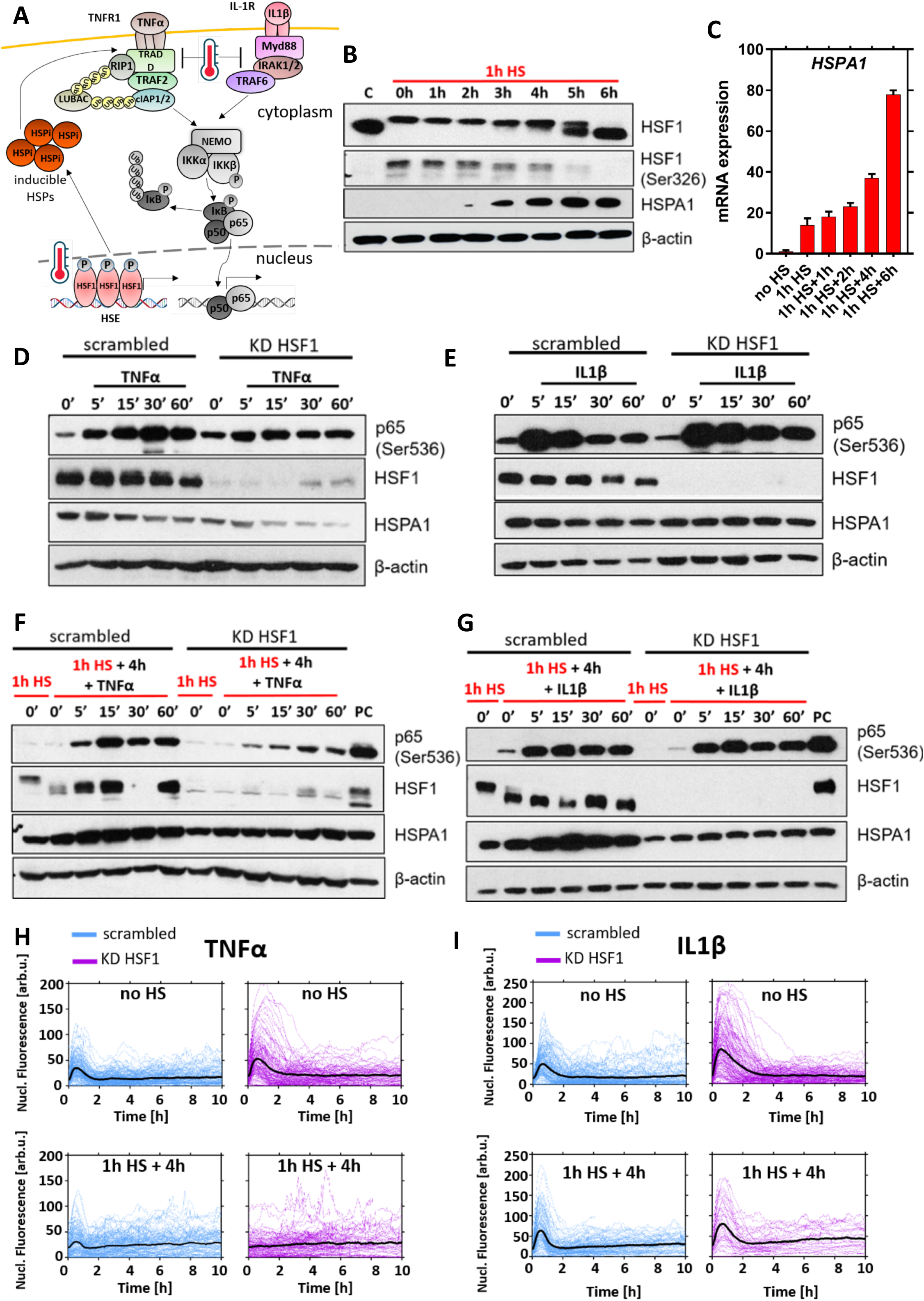
HSF1 regulates TNFα-induced NF-κB signalling recovery from HS. (**A**) Schematic representation of the HSR and NF-κB crosstalk. (**B**) Western blot analysis of HSF1, phosphorylated HSF1-Ser326 and HSP70A1 proteins level in MCF7 cells. Cells were either cultured in normal conditions, C, or subjected to 1h HS at 43°C and recovered for up to 6h. β-actin was used as a loading control. (**C**) Quantitative RT-PCR analysis of *HSPA1A* mRNA abundance in response to 1h HS at 43°C and recovery for up to 6h (normalised to the reference *GAPDH* gene). Shown are fold-changes (mean ± SDs of three replicate experiments) with respect to expression at 37°C (no HS control). (**D**) Effect of HSF1 knock-down on TNFα-induced response in MCF7 cells. Cells cultured in normal conditions were treated with scrambled siRNA control (scrambled) and HSF1 siRNA (KD HSF1) and stimulated with TNFα for indicated times (in min). Shown are the levels of p65-Ser536, HSF1 and HSPA1 assayed via Western blotting of the whole cell lysates. Shown also are untreated controls (0’) and β-actin loading control. (**E**) Effect of HSF1 knock-down on IL1β-induced response in MCF7 cells cultured and stimulated as in D. (**F**) Effect of HSF1 knock-down on TNFα-induced response following 4h HS recovery in MCF7 cells. Cells treated with scrambled siRNA control (scrambled) and HSF1 siRNA (KD HSF1) were subjected to 1h HS at 43°C and recovered for 4 h, then stimulated with TNFα for indicated times (in min). Shown are the levels of p65-Ser536, HSF1 and HSPA1 assayed via Western blotting of the whole cell lysates. Shown also are untreated controls (0’) and β-actin loading control. (**G**) Effect of HSF1 knock-down on IL1β-induced response following 4h HS recovery in MCF7 cells cultured and stimulated as in F. (**H**) Effect of HSF1 knock-down on TNFα-induced NF-κB response in MCF7 cells stably expressing p65-EGFP. Cells treated with scrambled siRNA control (scrambled) and HSF1 siRNA (KD HSF1) were either cultured in normal conditions (37°C, no HS) or subjected to 1h HS at 43°C and recovered for 4 hours (1h HS + 4h). Individual single cell nuclear p65-EGFP trajectories (n = 202, 127, 146, 98 respectively, in aribirary fluorescene units) are depicted with colour lines; population average – with a black line. (**I**) Effect of HSF1 knock-down on IL1β-induced NF-κB response in MCF7 cells stably expressing p65-EGFP. Cells were cultured and stimulated as in H (n = 153, 140, 113, 42 respectively). For heat maps and number of responding cells see Fig. S2.

The observed kinetics of HSPA1 protein accumulation coincided with the timing of the TNFα-induced NF-κB signalling recovery after HS (Fig. 1). We therefore sought to directly establish if HSF1 feedback controls these responses. For this purpose, we used siRNA to selectively knock-down the expression of HSF1 in MCF7 cells. The NF-κB activation by TNFα and IL1β was first confirmed in immunoblotting experiments both in cells treated with HSF1-specific or scrambled siRNA, which also confirmed a knock-down of HSF1 expression (Fig. 3D, E). Subsequently, we exposed cells to 1h HS and recovered for 4h before cytokine treatment, a condition which showed differential responses after TNFα and IL1β treatment. In cells treated with scrambled siRNA, we observed the recovery of the NF-κB signalling (visible as Ser536 p65 phosphorylation) (Fig. 3F, G) which was essentially the same as in wild type cells (Figs. 1B, 2C). Interestingly, HSF1 knock-down had no influence on the recovery of the NF-κB signalling in response to IL1β stimulation, while the responses to TNFα stimulation did not recover. As under normal conditions, an almost complete HSF1 knock-down was observed in these experiments and HSPA1 was up-regulated after HS only in cells treated with scrambled siRNA. Further microscopy analyses confirmed that under normal conditions, cells treated with HSF1-specific or scrambled siRNA showed robust p65-EGFP translocation to the nucleus following both TNFα and IL1β stimulation (Fig. 3H, I, top rows). After HS, markedly reduced p65-EGFP nuclear translocation was observed in cells treated with siRNA specific for HSF1 and stimulated with TNFα but not with IL1β (Fig. 3H, I, bottom rows; see Fig. S2 for heat maps and fraction of responding cells), which was consistent with differential Ser536 p65 phosphorylation. Overall, this data demonstrates that HSF1-dependent feedback regulates TNFα, but not IL1β-mediated NF-κB responses following HS.

### HS-mediated NF-κB responses are conferred via IKK signalosome

The attenuation of IKK activity via temperature-dependent denaturation and loss of solubility is thought to be critical for the NF-κB responses post HS (Lee et al., 2004; Pittet et al., 2005). We previously showed that exposure of human osteosarcoma cells to 1h 43°C HS resulted in depletion of soluble IKKα and IKKβ levels, effectively limiting the amount of IKK (and thus NF-κB) that can be activated by the cytokine stimulation (Kardynska et al., 2018). The cellular adaptation to HS requires restoration of IKK signalling. However, the kinetics of recovery and the relationship with HSR remain unexplored. Based on our findings that HSF1 is involved in the stimulus-specific NF-κB recovery post HS we developed a dynamical mathematical model of the HSR and NF-κB cross-talk to investigate these mechanisms more quantitatively.

We considered a simplified structure of the HSR pathway (Fig. 4A), which contains HSF1 and two HSP species: inducible (HSPi), transcription of which is strictly HSF1 dependent, and constitutive (HSPc) (Morimoto, 1993). Following previous work, we made a simplifying assumption that in resting cells HSF1 monomers are held in an inactive state via an association with HSPi, while HSPc acts as a generic chaperone for other proteins (Rybinski et al., 2013; Szymanska and Zylicz, 2009; Zheng et al., 2016). Temperature stimulation results in redistribution of HSPi (and HSPc) from the HSF1 complex, then HSF1 forms trimers and activates transcription of HSPi, creating a regulatory feedback loop that restores proteome homeostasis and eventually inhibits HSF1 activity (Richter et al., 2010). In addition, we utilised our existing model of the IKK-NF-κB signalosome (Adamson et al., 2016) involving TNFα and IL1β transduction pathways, each comprising a cognate receptor and upstream kinase (IKKK, Inhibitory κB kinase kinase) that in parallel regulate IKK activity (Fig. 4B). As in previous models (Adamson et al., 2016; Werner et al., 2008), cytokine-specific IKKKs denote generic IKK kinases, and simplistically represent complex and not fully elucidated signal transduction networks (DeFelice et al., 2019; Heyninck and Beyaert, 1999; Shembade et al., 2010). Subsequently, we considered different crosstalk mechanisms that could recapitulate the kinetics of cytokine-specific NF-κB responses after HS (for details of the mathematical modelling, see the Materials and methods section). Assuming that HS exposure results in ubiquitous damage of the proteins involved in the IKK signalosome, our data (Fig. 3) demonstrated that (1) the recovery of IL1β signalling was independent from the HSF1-mediated response, potentially mediated via action of constitutive HSPs; (2) the recovery of TNFα signalling depended on the inducible HSF1-HSPi response; which (3) was mediated via the signal-specific pathways upstream of IKK kinase (IKKK_TNF_). These mechanisms, when implemented in the combined cross-talk model, were able to very closely recapitulate the observed NF-κB signalling responses to TNFα and IL1β stimulation, in wild type and HSF1 knock-down cells (Fig. 4C, see also Tables S3 and S4 for model equations and fitted parameter values of the IKK-HSP interaction).

**Figure 4.**
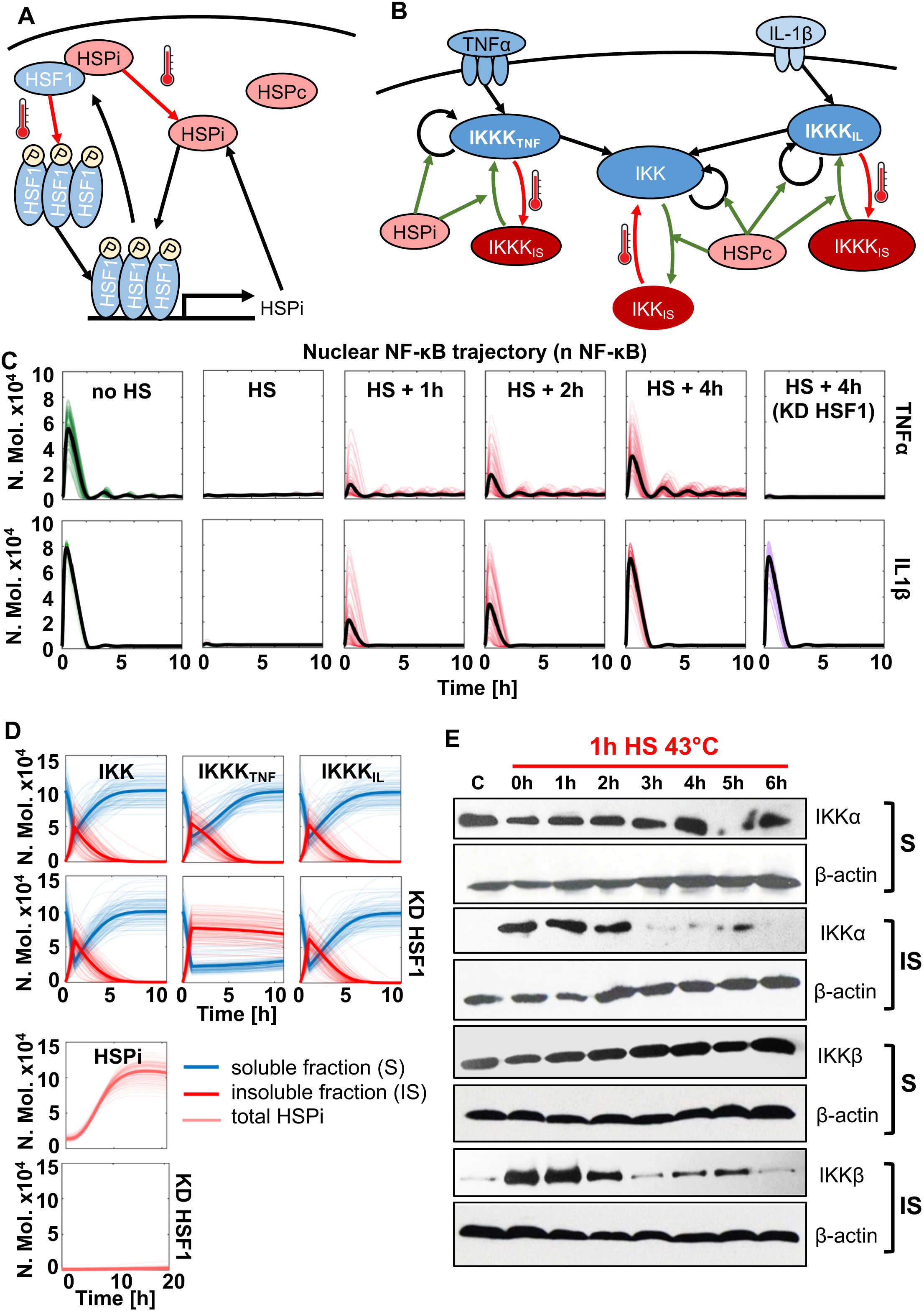
Mathematical model recapitulates HSR and NF-κB interaction via IKK signalosome. (**A**) Schematic representation of HSF1 signalling system. Red arrows indicate HS dependent regulation. (**B**) Schematic diagram of NF-κB and HS pathway crosstalk. In red is proposed temperature-dependent protein denaturation, in green interactions with HSP proteins. (**C**) Model simulations of the NF-κB-HSR crosstalk: wild type and HSF1 knock-down cells (KD HSF1) treated with TNFα (top) and IL1β (bottom) after different recovery times from HS (as indicated). Shown are sample 100 time-courses of nuclear NF-κB levels (coloured lines) and average nuclear NF-κB levels (in black), calculated from 1,000 single cell model simulations (in number of molecules). (**D**) Kinetics of the IKK signalosome. Simulations were performed for 1h HS at 43°C followed by 10h (IKK and IKKK) or 20h (HSPi) recovery time. Shown are time courses of simulated IKK, IKKK and HSPi levels in wild type cells (top) or cells with the HSF1 knock-down (KD HSF1) in number of molecules. (**E**) Western blot analysis of soluble (S) and insoluble (IS) IKKα and IKKβ proteins level in MCF7 cells. Cells were either cultured under normal conditions, C, or subjected to 1h HS at 43°C and/or recovered for 1-6 hours. β-actin was used as a loading control.

In the mathematical model, NF-κB behaviour is dictated by the changing levels of IKK activity (Werner et al., 2005). Simulations suggest that peak of denatured (insoluble) IKK occurs immediately after HS (at 1h), then it fully recovers to intact (soluble) forms following 4h post HS (Fig. 4D). The IL1β-specific IKK kinase (IKKK_IL_) recovers with similar kinetics. In contrast, TNFα-specific IKK kinase (IKKK_TNF_) recovers more slowly, which dictates the overall NF-κB signalling recovery post HS. In order to verify these predictions, we measured by immunoblotting the levels of soluble (intact) and insoluble (denatured) levels of IKKα and IKKβ subunits in MCF7 cells exposed to 1h HS and recovered for up to 6h (Fig. 4E). In lysates from cells cultured under normal conditions, IKKα and IKKβ were detected only in the soluble form. 1h HS exposure resulted in transition to insoluble form, evident of temperature-induced denaturation. The amount of insoluble IKKα and IKKβ fractions remained high at 2 h after HS exposure, but effectively returned to pre-HS state at 3h after exposure in agreement with model simulations. In addition, we investigated whether recovery after HS might depend on the cognate receptor availability and internalisation (Schneider-Brachert et al., 2004). Cells were stimulated with IL1β (and separately TNFα), fluorescently labelled with fluorescein isothiocyanate (FITC) and assayed under the microscope. In agreement with our modelling assumptions, no difference in the binding and internalisation of either cytokine was detected in cells exposed to HS in comparison to normal cell culture (Fig. S3). Overall, these analyses confirm that cytokine-specific IKK signalosome responses modify NF-κB signalling to 43°C HS by different mechanisms only partially depending on inducible HSF1 responses.

### NF-κB sensitivity to HS temperatures are cytokine specific

Mammalian cells experience a wide range of temperatures from physiological core body temperature and fever (<40°C) to heat stress used in clinical hyperthermia treatment (up to 45°C) (Maluta and Kolff, 2015). Having established the critical link between HSR and the NF-κB systems, we wanted to understand their sensitivities to a range of relevant temperatures. First, we assayed IKK solubility (Fig. S4A). The 1h exposure to different temperatures from a 38-43°C range resulted in increased insoluble IKKα and IKKβ levels in cell lysates (in comparison to cells cultured at 37°C). There was also visible an additional decrease in soluble IKK between 42 and 43°C. In addition, we used live-cell images of MCF7 cells expressing HSF1-dsRed to measure the HSR activation by examining the redistribution of the fusion protein into nuclear stress granules (Jolly et al., 1999) (Fig. S4B). We found a significant increase in the number of HSF1-dsRed granules at 41°C, compared to lower temperatures and the 37°C control (Fig. S4C). There was also a significant increase in number of HSF1 granules between 41 and 42°C, but no further increase at 43°C, which matched the temperature-dependent shift of the total HSF1 protein in the immunoblotting assay (Fig. S4D). Overall these data suggest temperature modulation over the 38-43°C range, with HSF1 response activated above 40°C. Our mathematical model was subsequently extended to incorporate the temperature-dependent IKK and HSF1 behaviour (Fig. S5). For simplicity, we assumed denaturation of IKK followed a nonlinear temperature-depended function and receptor-specific IKKKs shared the same characteristics (see Tables S3 and S4 for model description). Under these assumptions, 1h exposure to higher temperatures resulted in the gradual decrease of the soluble IKK and cytokine-specific IKKK levels (Fig. S5A). In turn, the graded protein denaturation resulted in a step-like HSF1 activation as a consequence of HSF1 redistribution form the HSP-HSF1 complex, in agreement with the data (Fig. S5B). As such, the model suggested that the level of IKK denaturation was closely linked to the level of HSF1 activation in the system (Fig. S5C).

Subsequently, using the developed model we performed comprehensive simulations to understand NF-κB attenuation and recovery following different temperature exposures (Figs 5A, B, S6 and S7). We found that in the case of the TNFα treatment, the model exhibited NF-κB inhibition following exposure to 41°C or higher temperatures (compared to cells cultured under normal condition). Immediately after 1h exposure to 41°C, the first peak nuclear NF-κB amplitude was 56% of that in control cells, while exposure to 42°C showed further inhibition to 31% (Fig. 5C). In contrast, IL1β-induced NF-κB signalling was predicted to be less sensitive to temperature changes; no inhibition was observed at 41°C, and only a partial inhibition was observed after 1h exposure to 42°C HS (Fig. 5D). In order to validate these predictions, we performed live-cell imaging studies at the critical 41°C temperature. In agreement with modelling, we found an inhibition of TNFα-induced responses in cells stimulated immediately after 1h 41°C exposure (i.e. significant reduction of AUC and peak amplitude as well as increased time to first response, compared to cells cultured under normal condition, Fig. 5E, F). As predicted, responses returned to pre-stimulation steady state in cells stimulated after 4h recovery. Also, we found no inhibition IL1β-mediated NF-κB responses by 41°C (in particular immediately after the exposure), validating the prediction of stimulus-specific temperature sensitivity (Fig. 5G, H). In the mathematical model, the increased sensitivity of TNFα-induced responses to HS was due to a lower NF-κB amplitude and thus IKK activity (in comparison to IL1β), which in turn was more affected by the level of protein denaturation at a given temperature (Figs S5D, S6B and S7B). Systematic sensitivity analyses confirmed that the kinetic parameters associated with the IKK module (but not IκBα feedback) control differential NF-κB responses to temperature changes (Fig. S8). This analysis also revealed a role in temperature regulation for the A20 protein feedback acting via IKK on temperature sensitivity in line with previous reports (Harper et al., 2018).

**Figure 5.**
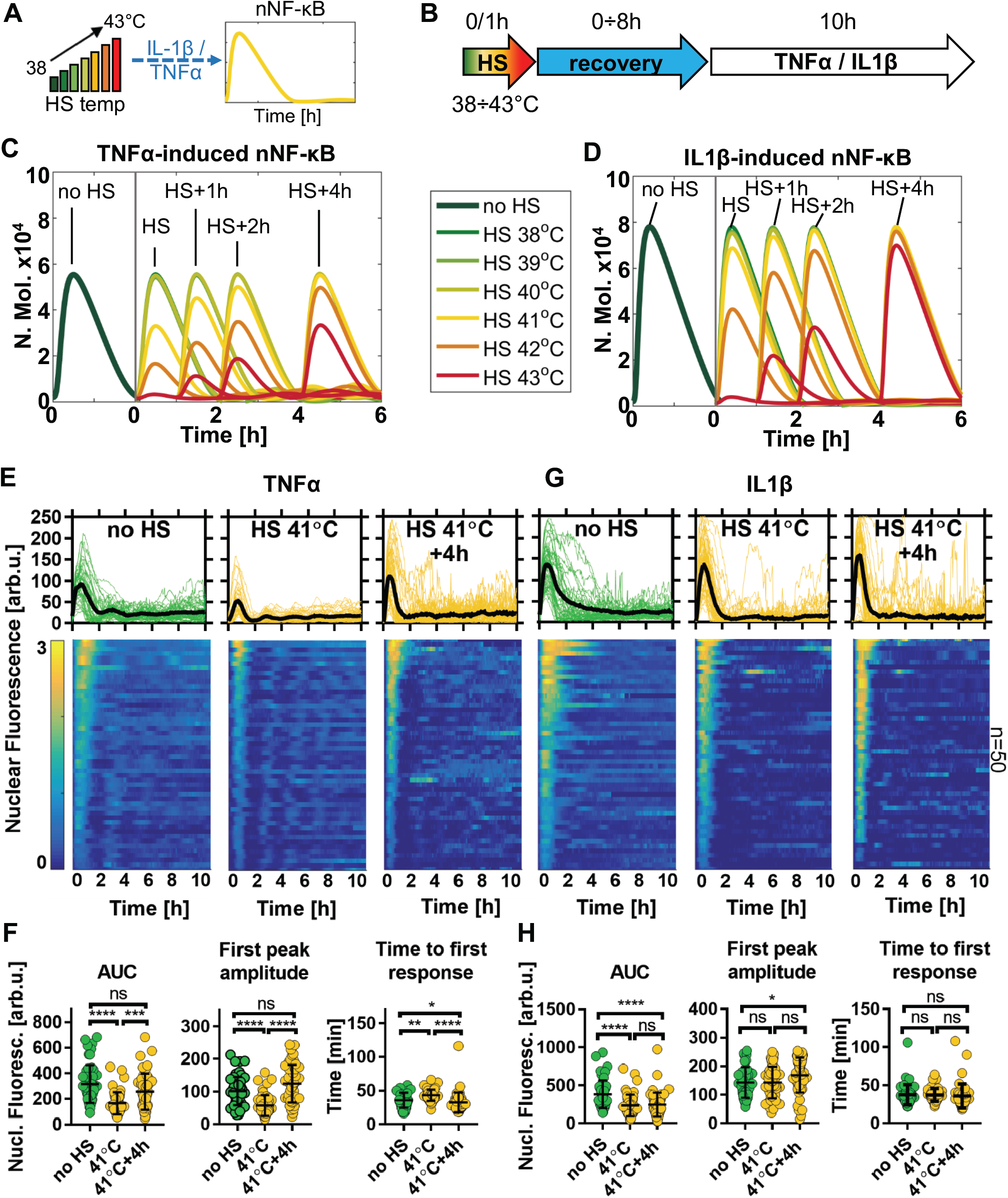
NF-κB responses exhibit cytokine-specific temperature sensitivity. (**A**) Schematic representation of the differential temperature treatment. (**B**) Schematic representation of the HS treatment protocol: cells exposed to 1h 38-43°C HS range and subjected to cytokine stimulation following different recovery time. (**C**) Model simulations of the TNFα-induced NF-κB responses following 1h temperature exposure and different recovery times. Shown are average nuclear NF-κB trajectories (based on 1,000 simulated cells, in number of molecules) for the 38-43°C temperature range and normal 37°C conditions, as indicated on the graph. (**D**) Model simulations of the IL1β-induced responses as described in C. (**E**) Nuclear NF-κB trajectories in MCF7 cells stably expressing p65-EGFP in response to TNFα stimulation. Cells were cytokine treated in normal conditions (left, data from Fig. 1), immediately after 1h 41°C exposure (middle), or after 4h recovery (right). Top: individual single cell nuclear NF-κB trajectories (n = 50 per condition, in aribirary fluorescene units) depicted with coloured lines (green 37°C, yellow 41°C); population average – depicted with a black line. Bottom: heat maps of trajectories normalized across all conditions in E and F (represented on a 0-3 scale). (**F**) Characteristics of TNFα-induced responses from E. From the left: distribution of area under the curve (AUC), first peak amplitude, and time to first response. Individual cell data are depicted with circles (with mean ± SD per condition). Kruskal-Wallis one-way ANOVA with Dunn’s multiple comparisons test was used to assess differences between groups (*p<0.05, **p<0.01, ***p<0.001, ****p<0.0001, ns – not significant). (**G**) Nuclear NF-κB trajectories in MCF7 cells stably expressing p65-EGFP following treatment with IL1β, represented as in E. p65-EGFP trajectories of cells cultured in normal conditions taken from Fig. 2. (**H**) Characteristics of IL1β-induced responses from G, data are represented as in F.

### Timing of HSP induction renders adaptation to repeated temperature stress

Overall, our analyses of cells exposed to elevated temperatures demonstrate that the HSF1-NF-κB crosstalk enables cytokine specific responses with differential temperature sensitivity. However, cells are also known to be able to adapt to repeated temperature challenges (Kampinga, 1993). This so-called thermotolerance effect is thought to depend on HSP accumulation (Lee et al., 2004; Pittet et al., 2005), which once induced may prevent IKK (and general proteome) damage to subsequent temperature exposures (Fig. 6A). Our data demonstrated that inducible HSPA1 accumulated in the system after 2-3h following exposure to 43°C (Fig. 3B), while simulations suggested accumulation over the 41-43°C range (Fig. S5E). We therefore used our crosstalk model to simulate NF-κB system responses following exposures to repeated temperature treatments at different time intervals ranging from 2 to 8h (Fig. 6B). We found that in the case of TNFα stimulation, exposure to elevated temperatures at 2h time interval resulted in NF-κB response inhibition in the 41-43°C range (Fig. S9A), when compared to a single HS exposure (Fig. S6). Thermotolerance, i.e. lack of NF-κB inhibition in cells exposed to repeated temperature treatment, was predicted to occur as early as 4h after the initial exposure (Fig. 6C). In contrast, IL1β-induced responses were predicted to be affected only at 2h exposure interval and only at 42 and 43°C temperatures (Fig. S9B), while responses at 4h were fully adapted (Fig. 6D). As before we performed live-cell imaging studies to validate these predictions. We found that at the critical 43°C temperature, cells treated with TNFα were adapted to the second temperature treatment (after 4h recovery from first HS). In these cases, the p65-EGFP nuclear amplitude could not be distinguished from that in cells treated 4h after a single HS exposure (Fig. 6E, F. Of note, some cells exhibited delayed response times (comparing to a single HS treatment or in fact cells treated under normal conditions) highlighting increased heterogeneity of the responses. Also, as predicted by our mathematical model, IL1β-induced responses following repeated 43°C exposure were fully adapted, i.e. p65-EGFP amplitude was similar to that of a single HS treatment as well as of cells treated in normal conditions (Fig. 6G, H). In the mathematical model, the apparent HSP accumulation prevented IKK signalosome damage to the second HS exposure facilitating a normal IL1β-induced response (Fig. S9A, B). In agreement, immunoblotting analyses demonstrated lack of IKKα and IKKβ denaturation following exposure to repeated HS after 3h recovery (Fig. S9C). Overall, these analyses validate the predictive power of our mathematical model and demonstrate that via the HSR feedback the NF-κB system may rapidly adapt to repeated temperature treatment.

**Figure 6.**
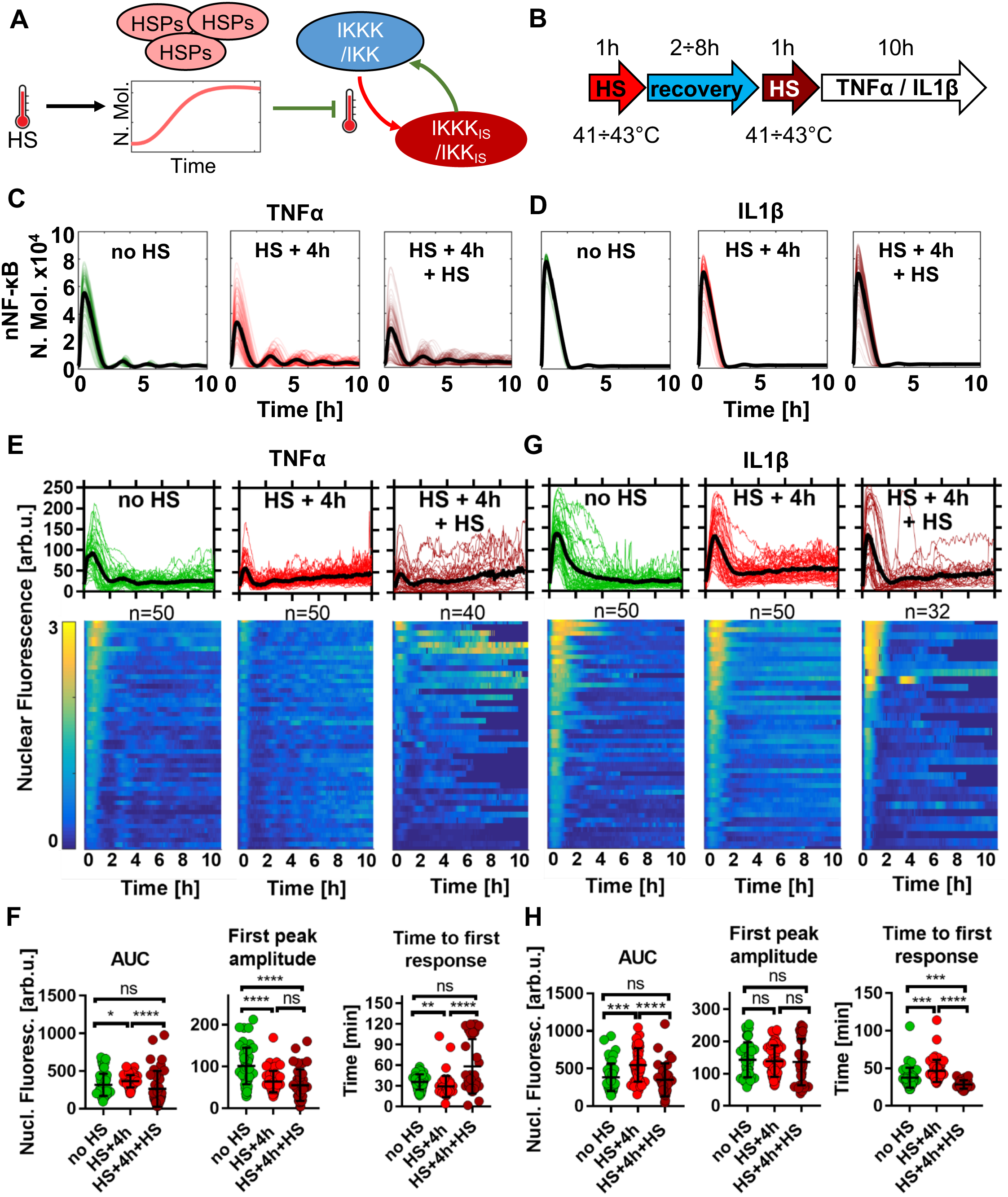
NF-κB adaptation to repeated temperature exposure. (**A**) Schematic representation of the thermotolerance: accumulation of inducible HSPs prevents IKK denaturation (in green) to repeated HS exposure (in red). (**B**) Schematic representation of the repeated HS treatment protocol: cells exposed to two 1h HS at indicated interval and subjected to cytokine stimulation. (**C**) Model simulation of TNFα-induced NF-κB responses following different HS protocols. Shown are sample 100 trajectories of single cell (and average responses based on 1,000 simulated cells, in number of molecules, in black) treated in normal conditions (left, in green), 4h after a single 1h 43°C HS exposure (middle, in red) and immediately after second 1h 43°C HS exposure (right, in burgundy). (**D**) Model simulation of IL1β-induced NF-κB responses following different HS protocols, as in C. (**E**) Nuclear NF-κB trajectories in MCF7 cells stably expressing p65-EGFP in response to TNFα stimulation. Cells are either treated in normal conditions (left, data from Fig. 1), 4h after a single 1h 43°C exposure (middle, data from Fig. 1) or immediately after second 1h 43°C HS exposure (right). Top: individual single cell nuclear NF-κB trajectories (n = 50, 50 and 40 per condition, respectively, in aribirary fluorescene units) depicted with coloured lines; population average – depicted with a black line. Bottom: heat maps of trajectories normalized across all conditions in E and F (represented on a 0-3 scale). (**F**) Characteristics of TNFα-induced responses from E. From the left: distribution of area under the curve (AUC), first peak amplitude, and time to first response. Individual cell data are depicted with circles (with mean ± SD per condition). Kruskal-Wallis one-way ANOVA with Dunn’s multiple comparisons test was used to assess differences between groups (*p<0.05, **p<0.01, ***p<0.001, ****p<0.0001, ns – not significant). (**G**) Nuclear NF-κB trajectories in MCF7 cells stably expressing p65-EGFP following treatment with IL1β, represented as in E. p65-EGFP trajectories of cells cultured in normal conditions or exposed to IL1β after 4h recovery from 1h 43°C are taken from Fig. 2. (**H**) Characteristics of IL1β-induced responses from G, data are represented as in F.

## Discussion

Temperature perturbs key cellular processes and cell function, in particular, involved in proliferation and inflammatory signalling (Ritossa, 1996). Thus, the ability to sense temperature changes to restore homeostasis is one of the most fundamental cellular response systems (Richter et al., 2010). Here we provide a new quantitative understanding of the dynamical crosstalk mechanisms involved in the regulation of the HSR and NF-κB networks using live single-cell microscopy and mathematical modelling approaches. We demonstrate that the kinetics of the NF-κB system following temperature stress is cytokine specific and may exhibit rapid adaptation to temperature changes. In agreement with previous findings, the exposure of breast adenocarcinoma cells to 43°C HS resulted in the attenuation of the immediate NF-κB signalling and gene expression response to TNFα and IL1β stimulation. However, while the IL1β-induced responses return to the normal level within 4h after HS exposure, the recovery following TNFα-mediated responses is delayed. TNFα and IL1β are two cytokines that confer related but distinct pro-survival and pro-inflammatory functions. Their expression is elevated in the local or inflamed tissue, e.g. in breast cancer (Wee et al., 2015), and their signalling, in the context of elevated temperature is important for physiological and clinical cellular responses (Perkins, 2012). Previous studies have focused solely on TNFα-mediated NF-κB signalling (Esquivel-Velazquez et al., 2015; Janus et al., 2015; Kardynska et al., 2018; Lee et al., 2004; Wong et al., 1997b), or examined a combined effect of cytokine mixes (Pittet et al., 2005). Here we show that the stimulus-specificity is rendered via HSR differentially controlling TNFα and IL1β signal transduction pathways. We argue that these data essentially reveal that (1) the recovery of IL1β signalling is independent of the inducible HSF1 response, potentially mediated via action of constitutive HSPs; (2) the recovery of TNFα signalling depends on the inducible HSF1-HSPi response; which (3) is mediated via the signal-specific pathways upstream of the IKK signalosome. Moreover, we demonstrate that individual cytokines essentially exhibit different temperature sensitivity and adaptation to repeated HS when exposed to a 37-43°C temperature range. Specifically, IL1β-mediated NF-kB responses are more robust to temperature changes in comparison to those induced by TNFα treatment.

Several lines of evidence highlight the modulation of IKK signalosome and the concurrent HSF1-HSP feedback activation as the key regulators of responses to elevated temperature (Esquivel-Velazquez et al., 2015; Janus et al., 2015; Kardynska et al., 2018; Lee et al., 2004; Lee et al., 2005; Morimoto, 1998; Perkins, 2004; Wong et al., 1997b). It is thought that physiological temperatures (<40°C) modulate IKK and consequently fine-tune NF-κB responses (oscillations) via the timing of the A20 negative feedback (Harper et al., 2018). In contrast, responses to extreme temperatures (>40°C) have been shown to involve direct attenuation of IKK activity via temperature-dependent denaturation and loss of solubility (Kardynska et al., 2018; Lee et al., 2004; Pittet et al., 2005). Consequently, the temporal adaptation to HS requires restoration of IKK and upstream transduction pathways (Salminen et al., 2008). Consistently with this idea, here we observe the temperature-dependent gradual loss of IKKα and IKKβ solubility over a 38-43°C range, levels of which return to the resting steady state within 6h post HS. We fully expect that several additional mechanisms might contribute to the observed behaviour (Kardynska et al., 2018). These might include modulation of IKK kinase activity (in addtion to protein level) (Shembade et al., 2010), inhibition of NF-κB transport (Furuta et al., 2004) or level (Sheppard et al., 2014), transcriptional co-regulation of NF-κB-dependent genes by HSF1 (Janus et al., 2015) and in particular, diverse action of specific HSP molecules (Chen et al., 2002). When it comes to signal specificity upstream of IKK and delayed recovery of TNFα-induced responses, each transduction pathway acts in parallel. TNFα signalling involves TRAF2/RIP, while IL1β involves TRAF6/IRAK-mediated pathways via enzymatic interactions with A20 (Heyninck and Beyaert, 1999; Shembade et al., 2010). The mechanisms acting in their differential HSR control remains to be elucidated. One interaction that might be important in HSF1-dependent modulation of TNFα signalling is the apparent temporal sequestration of TRAF2 adaptors into stress granules following HS (Kim et al., 2005), and thus the requirement for their recovery. In this work, using live single-cell microscopy we have the ability to monitor the heterogeneity of NF-κB signalling responses (Paszek et al., 2010). While responses appear to be more homogenous than those reported in the previous work using human osteosarcoma cells (Kardynska et al., 2018), we observe that TNFα-induced oscillatory patterns become more robust following recovery from HS in comparison cells treated under normal condition. As such, this is consistent with the idea that inducible HSPs (given their long half-lives) (Lee et al., 2005) might be able to influence NF-κB responses and potentially target gene expression over prolonged periods of time.

The ultimate goal of mathematical modelling is to interpret data and make biological predictions (Kirk et al., 2015). Here we developed and validated a dynamical mathematical model of NF-κB and HSR crosstalk, which combines previously published network structures (Adamson et al., 2016; Rybinski et al., 2013; Szymanska and Zylicz, 2009; Zheng et al., 2016). We proposed a crosstalk mechanism to recapitulate our original data on NF-κB p65 responses in wild type and HSF1 knock-down cells as well as measurements of the IKK denaturation and activation of the HSF1-HSP pathway for a 37-43°C temperature range. Here, we make simplifying assumptions, which essentially allow us to directly relate the level of IKK damage to the level HSF1 activation for a given temperature suggesting that the former is indicative of the overall proteome damage. Using our developed mathematical model, we systematically screened the NF-κB signalling responses to a single and repeated HS exposures for a range of temperatures and recovery times. We predicted critical conditions where the TNFα and IL1β-mediated responses exhibit differential sensitivity to temperature, or HSP-mediated thermotolerance. Subsequently, we performed additional imaging experiments to validate these predictions. These specifically demonstrate that the TNFα-induced NF-κB signalling responses are attenuated at 41°C, while the corresponding IL1β-induced signalling remains intact. In combination with sensitivity analyses this reinforces the idea that the IKK signalosome is a *bone fide* temperature sensor (Sengupta and Garrity, 2013), which effectively enables NF-κB signalling responses with cytokine specificity and differential temperature sensitivity. It will be important to understand the biophysical basis for the apparent temperature sensitivity of IKK subunits and molecules involved in signal transduction. Of note, we previously showed that the NF-κB p65 appears to be more stable than IKK molecules and does not undergo denaturation with 1h 43°C HS (Kardynska et al., 2018). Whether in general the signal transduction molecules are more temperature sensitive than other network components would be important to understand (Wallace et al., 2015).

The kinetic HSR and the NF-κB crosstalk is clinically relevant to hyperthermia, an emerging strategy to sensitise cancer cells to treatment (by chemotherapy or radiotherapy) with an artificial increase of tissue temperature (Wust et al., 2002). The efficacy of hyperthermia treatment is thought to critically depend on the timing between the HS exposure and treatment, in order to maximise the effect of proapoptotic proteome damage and minimise the effect of prosurvival induction of thermotolerance (Rybinski et al., 2013). Current treatment protocols utilise a wide time window, where HS in the range of 38-45°C is applied for up to 24h before and after treatment (Maluta and Kolff, 2015). Our analyses demonstrate that the time windows rendering cells sensitive to treatment may be short even at extreme temperatures, i.e. up to 4h at 43°C, completely absent at temperatures below 41°C for IL1β, while post HS cells may achieve more robust responses. While cells exhibit more sensitivity to other cytokines like TNFα stimulation, its effect on the NF-κB system activation is likely to be compensated by IL1β in the local tissue. In general, we expect that the NF-κB system (or in fact the combined action of other systems, including MAP kinase (Rouse et al., 1994) or p53 tumour suppressor (Alexandrova and Marchenko, 2015)) in individual cell types might exhibit different sensitivities and recovery kinetics following temperature exposure. For example, we previously showed that TNFα stimulation in human osteosarcoma cells resulted in “all-or-nothing” NF-κB responses following HS (Kardynska et al., 2018), while tissue-level architecture might impose additional spatial constraints (Bagnall et al., 2018). We suggest that further efforts should combine dynamical modelling with cell fate, in order to better understand the relationship between temperature and NF-κB as well as cell proliferation and apoptosis in the more relevant cancer or inflammatory context. However, it is clear that more efficient hyperthermia treatment protocols require better quantitative understanding of the underlaying processes.

## Materials and methods

### Cell culture and reagents

Experiments were performed using the human MCF7 adenocarcinoma cell line (purchased from ATCC^®^; cat. no. HTB-22™). Cells were cultured at 37°C in humidified 5% CO_2_ in DMEM/F12 medium (Gibco) supplemented with 10% (v/v) heat-inactivated foetal calf serum and routinely tested for mycoplasma contamination. For confocal and Western blotting experiments, HS response was induced by transferring cells into a water bath at 43°C (unless otherwise stated). After 1 hour of HS, cells were supplemented with fresh 37°C media. 10 ng/ml of human recombinant TNFα or IL1β (Calbiochem) was used to simulate NF-κB responses for the indicated time periods.

### Engineering of p65-EGFP and HSF1-dsRed stably transfected MCF7 cell line

The p65-EGFP sequence was re-cloned from p65-EGFP-N1 plasmid (Nelson et al., 2004) into pLNCX2 vector (Clontech) for expression under a CMV promoter (using *HindIII* and *Not1* restriction sites). HSF1 coding sequence was amplified by PCR on cDNA template, cloned in frame in dsRed-N1 plasmid and then HSF1-dsRed was re-cloned into pLNCX2 vector (using *AgeI* and *NotI* restriction sites). The Retroviral Gene Transfer protocol (Clontech) was followed for to obtain stable MCF7 line expressing the resulting p65-EGFP-pLNCX2 or HSF1-dsRed-pLNCX2 vector. In brief, RetroPack™ PT67 packing cells were transfected with p65-EGFP-pLNCX2 or HSF1-dsRed-pLNCX2 the vector using TurboFect™ (Thermo Scientific), then cells were selected with G418 geneticin sulphate (Gibco), and the virus-containing medium was collected after one week of culture. MCF7 cells were exposed to the virus-containing medium and transduced cells were sorted based on EGFP or dsRed expression.

### Protein extraction and Western blotting

Cells were lysed in 1% NP-40, 0.5% sodium deoxycholate and 0.1% SDS in PBS, supplemented with Complete™ (Roche) protease and phosphatase inhibitor cocktail, centrifuged for 20 min at 14,000 rpm at 4°C. The supernatants, defined as a soluble fraction, were collected. Insoluble proteins, remaining in the pellets, were dissolved in a SDS sample buffer consisting of 25 mM Tris-HCl pH 6.8, 0.5% SDS, 2.5% glycerol, and 15% 2-mercaptoethanol and sonicated (30 times 30s). Protein concentration was determined by BCA assay (Thermo Scientific). Samples and ladder (Bio-Rad, #161-0375) were resolved on polyacrylamide gels and transferred to nitrocellulose membranes (Amersham), incubated 1h at room temperature in blocking buffer (5% (w/v) non-fat milk powder in TBS-T containing 0.25 M Tris–HCl pH 7.5, 0.1% Tween-20, 0.15 M NaCl), washed 3 times in TBS-T and incubated overnight with primary antibody (p65-S536, CST #3033; IKKα, CST #11930S; IKKβ, CST #2684S; HSF1, CST #4356; HSF1-S326 Abcam ab76076; HSPA1, Stressgen ADI-SPA-810-D; β-actin, Sigma-Aldrich A3854) at 1:1,000 dilution in blocking buffer. Membranes were washed 3 times in TBS-T and incubated with 1:1,000 HRP-conjugated secondary antibody for 1h at RT. Membranes were washed (3 × TBS-T) then incubated with Luminata Crescendo Western HRP Substrate (EMD Millipore Corp.) and the signal was detected by exposure to Carestream Kodak BioMax MR film (Sigma-Aldrich).

### HSF1 siRNA knock-down

MCF7 or p65-EGFP expressing cells were plated into 35mm culture dishes one day before transfection. Transfection mix was prepared using DharmaFECT 1 Transfection Reagent (GE Dharmacon) according to the manufacturer’s protocol. Each dish was transfected with 100nM of human HSF1 On-Target Plus siRNA or On-Target Plus non-targeting pool siRNA (both GE Dharmacon). Cells were cultured with transfection mix for 48h. After 48h the entire procedure was repeated. Cell medium was replaced with fresh one and fresh transfection mix. After a further 48h cells were used in appropriate experiments.

### Gene expression analysis

Total RNA was extracted from wild type MCF7 cells using the Roche High Pure RNA Isolation Kit. The mRNA concentration and purity was quantified using a NanoDrop spectrophotometer. The absorbance was measured at 260 and 280 nm. All mRNA were of sufficient purity with an A260/A280 ratio between 1.9-2.1 used in the reverse transcriptase reaction. SuperScript™ VILO™ kit (Invitrogen) was used for production of cDNA. For each sample, 2 μg of mRNA was used. Manufacturer’s protocol was followed. The cDNA was diluted 1:20 with RNase free water. Quantitative RT-PCR was performed using Roche LightCycler® 480 Instrument II. A total of 5 pM of forward and reverse primers, cDNA template was added to the 2x LightCycler® 480 SYBR Green I Master (Roche). Primers used in the analyses are listed in Table S1. Relative quantification was used to calculate the fold difference based on the threshold cycle (CT) value for each PCR reaction using 2^−ΔΔCT^ method. The target gene was normalised to the reference gene *GAPDH*, with no HS and NT control used as the calibrator.

### Confocal microscopy

Cells were plated onto 35 mm-glass-bottomed dishes (Greiner Bio-One) one day prior to the experiment and incubated on the microscope stage at 37°C in humidified 5% CO_2_. The Hoechst 33342 (Molecular Probes) staining was performed immediately before the experiment. Two Carl Zeiss confocal microscopes were used (LSM780, AxioObserver and LSM880 AxioObserver) with Plan-Apochromat 40x/1.4 Oil DIC M27 and Fluar 40x/1.30 M27 Oil objectives. The 488 nm (ATOF set at 4%) line from an argon ion laser was used to excite the p65-EGFP fusion protein and emitted light between 498-598 nm was detected through pinholes set to 5 µm. The 405 nm (ATOF set at 1%) line from a diode laser was used to excite Hoechst 33342 and emitted light 410-490 nm was detected. The 556 nm (ATOF set at 2%) line from a diode laser was used to excite the HSF1-dsRed fusion protein and emitted light 580-650 nm was detected. For the series of interrelated confocal experiments, the same microscope settings have been used. Image capture was performed using the Zeiss Zen 2010b or Zen2 software. Quantification of p65-EGFP nuclear fluorescence was performed using automated segmentation and tracking of Hoechst-labelled cell nuclei with Cell Tracker (version 0.6) (Shen et al., 2006) and in-house software. The data was exported as mean fluorescence intensity. Trajectories of the nuclear p65-EGFP were normalized across presented conditions and displayed as heat maps. HSF1-dsRed granule quantification was performed in CellProfiler with a modified speckle counting pipeline (Carpenter et al., 2006).

### Evaluation of TNFα and IL1β internalization

MCF7 cells were plated onto 4-compartment 35 mm-glass-bottomed imaging dishes (Greiner Bio-One) in culture medium one day prior to the experiment and incubated at 37°C in humidified 5% CO_2_ on the microscope stage. Cells were treated with human recombinant TNFα or IL1β biotin conjugate (1 µg/ml, Fluorokine, R&D Systems, Wiesbaden) diluted to 10 ng/ml in 20 µl of avidin-FITC (10 µg/ml) and made up to 50 µl with minimum essential medium. Carl Zeiss LSM880, AxioObserver confocal microscope with a Plan-Apochromat 40x/1.4 Oil DIC M27 objective was used with 488 nm excitation and 493-634nm emission signal detection. Image capture was performed using Zeiss Zen 2 software to take time lapse 13 deep Z stacks over 13µm with a 1 Airy unit pinhole diameter. Maximum intensity projections were used for image analysis.

### Statistical analyses

Statistical analyses were performed in GraphPad Prism 7.02. Normal distribution was assessed with D’Agostino-Pearson test. Nonparametric tests were applied for non-normal distribution data. Kruskal-Wallis one-way ANOVA with Dunn’s multiple comparisons was used for characteristics of single cell NF-κB responses. Differences in the percentage of responding cells were assessed with Chi-square test.

### Mathematical modelling

We considered simplified models of HS-induced HSPi protein accumulation (Rybinski et al., 2013; Szymanska and Zylicz, 2009; Zheng et al., 2016) combined with a previously published model of TNFα and IL1β-dependent NF-κB signalling (Adamson et al., 2016). Models were fitted to recapitulate: (1) HSF1 activation (via release from the HSP-HSF1 complex due to protein denaturation) and accumulation of HSPi proteins (via transcriptional regulation with a Hill coefficient n = 3 corresponding to HSF1 trimerization). (2) MCF7-specific NF-κB dynamics; namely, the dampening and low first peak amplitude of NF-κB oscillations in response to TNFα was recapitulated by a reduced IκBα transcript rate (Wang et al., 2012) and IKKK_TNF_ activation rate, while a single translocation in response to IL1β by a slower rate of IKKK_IL1_ cycling (in comparison to IKKK_TNF_). Subsequently, the HSR and NF-κB cross-talk was introduced in the combined model by assuming (1) temperature nonlinearly affects denaturation rate of IKK and IKKK kinases, and no other molecules in the system; (2) HSPs prevent denaturation of IKK and IKKK kinases (Kampinga, 1993); (3) Repair of the TNFα receptor-associated kinase (IKKK_TNF_) requires HSPi; (4) Repair of IKK and IL1β receptor-associated kinases (IKKK_IL1_) involves HSPc; (5) HSPs accelerate the transition between active and inactive kinase states. Crosstalk parameters were fitted to recapitulate single cell NF-κB responses (to TNFα and IL1β stimulation) in wild-type and HSF1 knock-down cells at different times after HS as well as data on IKK denaturation and HSF1 activation. The final mathematical model is presented, including 31 ordinary differential equations and 60 parameters that describe protein association/dissociation/degradation and mRNA transcription/translation using mass action kinetics (see Figs 4C, D and S5A, B for model fits, Table S3 and S4 for differential equations and parameter values). Model simulations were divided into three phases: (i) heat shock – 1h HS at 43°C (Fig. 4) and subsequently extended to 38-43°C range (Figs 5, S6 and S7), (ii) recovery time – up to 8 hours at 37°C, (iii) TNFα or IL1β stimulation for 10h at 37°C. Heat shock treatments were subsequently repeated to test the effect of thermotolerance (Fig. S9). In order to simulate heterogeneous cell responses, random initial numbers of molecules for IKK and IKKK, as well as HSPc and HSF1 were assumed. Initial conditions were drawn from the log-normal distribution with parameters µ = 11.5 and σ = 0.15 for IKK, IKKK, µ = 10.1 and σ = 0.15 for HSPc, and µ = 9.2 and σ = 0.15 for HSF1 molecules. In simulations with HSF1 knock-down, a 95% reduction in the amount of HSF1 molecules was assumed. Sensitivity analyses were performed by varying temperature (by 1°C) and parameter values (one parameter at a time over an 8-fold range comparing with the nominal value for a given cytokine transduction pathway), results were presented as heat maps (Fig. S8). Simulations were performed in MATLAB R2016a with ‘ode15s’ function.

## Acknowledments

We thank Michael White and other members of Systems Microscopy Centre in Manchester for discussions. This work has been supported by National Science Centre, Poland (https://ncn.gov.pl), grants 2016/23/B/ST6/03455 (AP, MKard), 2015/19/B/ST7/02984 (MK) and 2016/21/B/ST7/02241 (PP), Silesian University of Technology grants 02/010/BK19/0143 (JS), Biotechnology and Biological Sciences Research Council (http://www.bbsrc.ac.uk) grant BB/K003097/1 (AP, PP and MW). Publication supported by scholarship financed from Silesian University of Technology scholarship fund, contract number 825/RN2/RR4/2018 (MKard). AP was supported by Fundacja Jakuba Hrabiego Potockiego. Calculations were performed on the Ziemowit computational cluster (http://www.ziemowit.hpc.polsl.pl) created in the POIG.02.01.00-00-166/08 project (BIO-FARMA) and expanded in the POIG.02.03.01-00-040/13 project (Syscancer).

