## Supplementary Figures and Tables for "Heat shock response pathways regulate stimulus-specificity and sensitivity of NF-κB signalling to temperature stress"

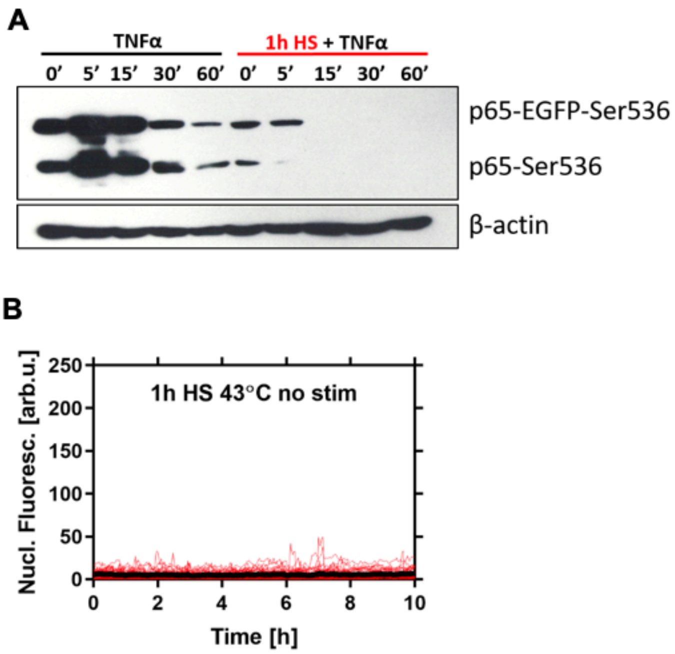


**Figure S1. Analysis of NF-κB signalling responses in MCF7 cells.** (**A**) Analysis of NF-κB p65-Ser536 phosphorylation in transformed cells. The level of p65-Ser536 phosphorylation was analyzed by Western blot in the whole MCF7 p65-EGFP cells lysates. Cells either cultured under normal conditions (37°C) or subjected to 1h HS at 43°C were treated with TNFα for indicated times. β-actin was used as a loading control. (**B**) Nuclear NF-κB trajectories in MCF7 cells stably expressing p65-EGFP after 1h 43°C HS treatment. Individual single cell trajectories (n = 50 per condition) are depicted with colour lines; population average is depicted with a black line.

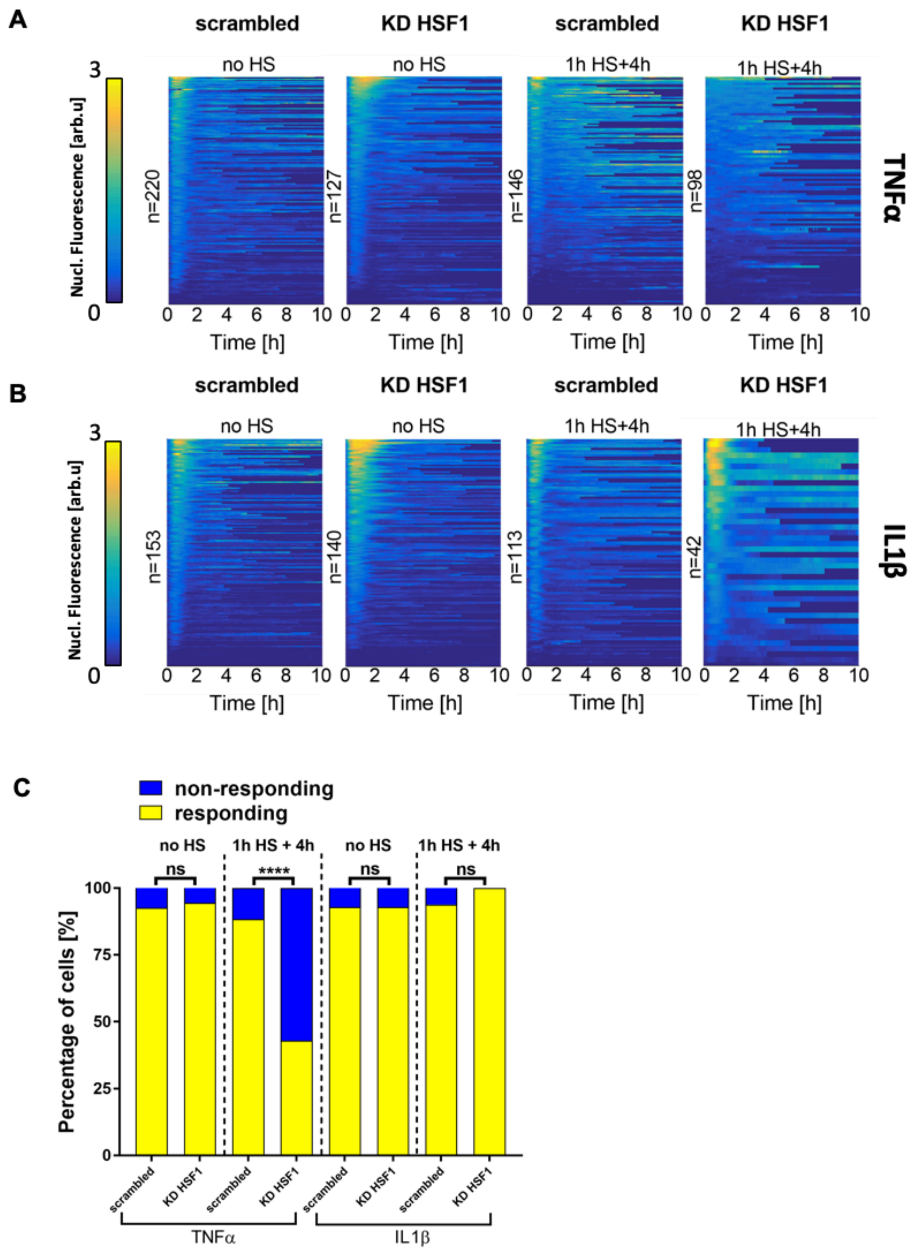


**Figure S2. Analysis of NF-κB responses in HSF1 knock-down cells.** (**A**) Heat maps of nuclear NF-κB trajectories in response to TNFα in MCF7 cells stably expressing p65-EGFP. Cells were treated with scrambled siRNA control (scrambled) or HSF1-specific siRNA (KD HSF1) and stimulated with the cytokine under normal conditions (37°C, no HS) or after 1h HS at 43°C and 4 hours recovery (1h HS + 4h). Heat maps of trajectories were normalized across all conditions (represented on a 0-3 scale). Individual single cell trajectories are shown. (**B**) Heat maps of nuclear NF-κB trajectories in response to IL1β in MCF7 cells stably expressing p65-EGFP. Cells were treated and data are presented as in A. (**C**) Percentage of cells responding (yellow) and non-responding (blue) to stimulation with TNFα or IL1β (from data shown in A and B). Statistical difference was assessed with Chi-square test (****$p < 0.0001$, ns – not significant).

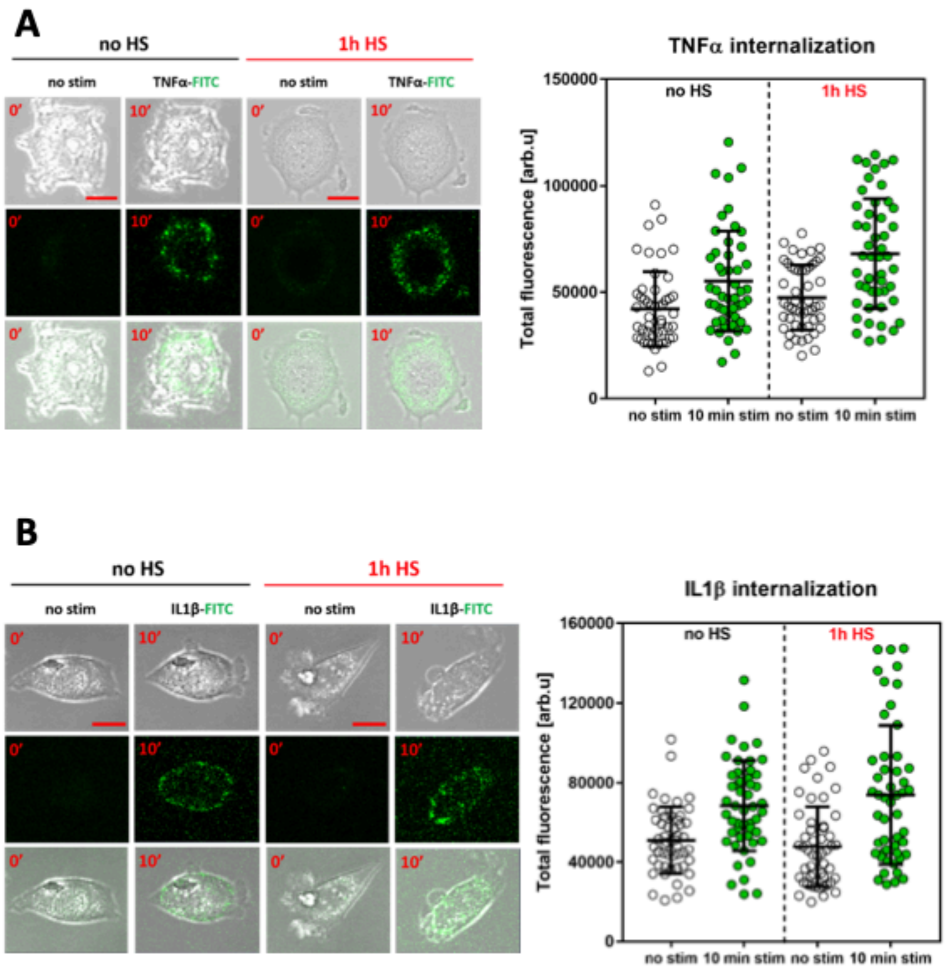


**Figure S3. Analysis of cytokine uptake after HS.** (**A**) Confocal microscopy images of representative MCF7 cells stimulated with fluorescently labelled TNFα. Cells were cultured under normal conditions (at 37°C, no HS) or exposed to 1h HS at 43°C prior to TNFα stimulation. FITC-conjugated TNFα was applied at 0 min and measured 10 min after stimulation. Top – bright field, middle – FITC, bottom – merged images. Scale bar, 10 µM. On the right: quantified individual cell fluorescent levels as well as mean ± SD per condition, based on three experimental replicates. (**B**) Confocal microscopy images of representative MCF7 cells stimulated with fluorescently labelled IL1β and quantified fluorescence levels (as in A).

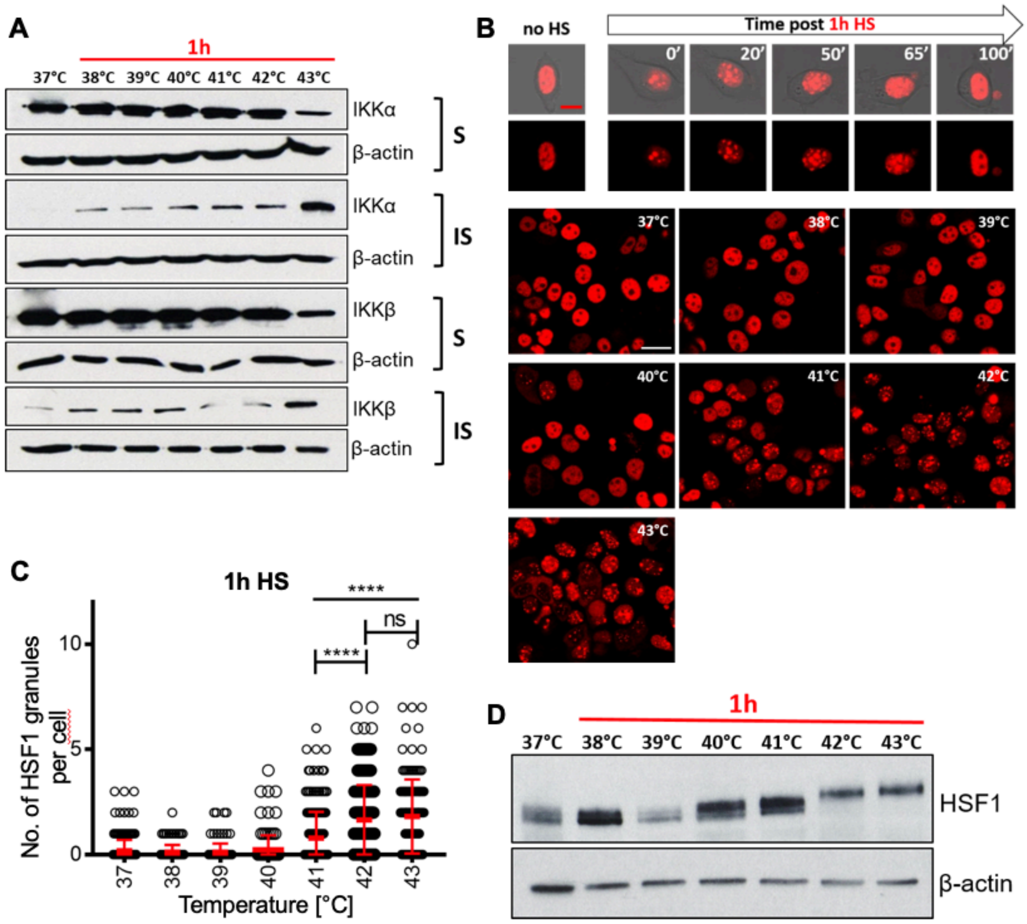


**Figure S4. Temperature sensitivity of the NF-κB and HSR signalling.** (**A**) Western blot analysis of soluble (S) and insoluble (IS) IKKα and IKKβ proteins level in MCF7 cells. Cells were either cultured under normal conditions, 37°C, or subjected to 1h temperature shift (38-43°C range, as indicated on the graph). β-actin was used as a loading control. (**B**) Temperature-sensitivity of HSF1 stress granule formation. Confocal microscopy images of representative MCF7 cells stably expressing HSF1-dsRed fusion protein. (Top) Cells cultured under normal conditions (at 37°C, no HS) or exposed to 1h HS at 43°C and imaged thereafter. Recovery time after HS displayed in minutes. Scale bar 5 µm. (Bottom) Cells assayed under normal conditions (at 37°C, no HS) or assayed following 1h HS at 38-43°C temperature range. Scale bar 10 µm. (**C**) Distribution of stress granules in MCF7 cells stably expressing HSF1-dsRed. Individual cell data as in B are depicted with circles (with mean ± SD per condition, of >117 cells per condition). Kruskal-Wallis one-way ANOVA with Dunn's multiple comparisons test was used to assess differences between groups (****p<0.0001, ns – not significant). (**D**) Western blot analysis of the total HSF1 protein level in MCF7 cells. Cells were either cultured in normal conditions, C, or subjected to 1h temperature stress in the 38-43°C range. β-actin was used as a loading control. Shift of the HSF1 band indicates activation.

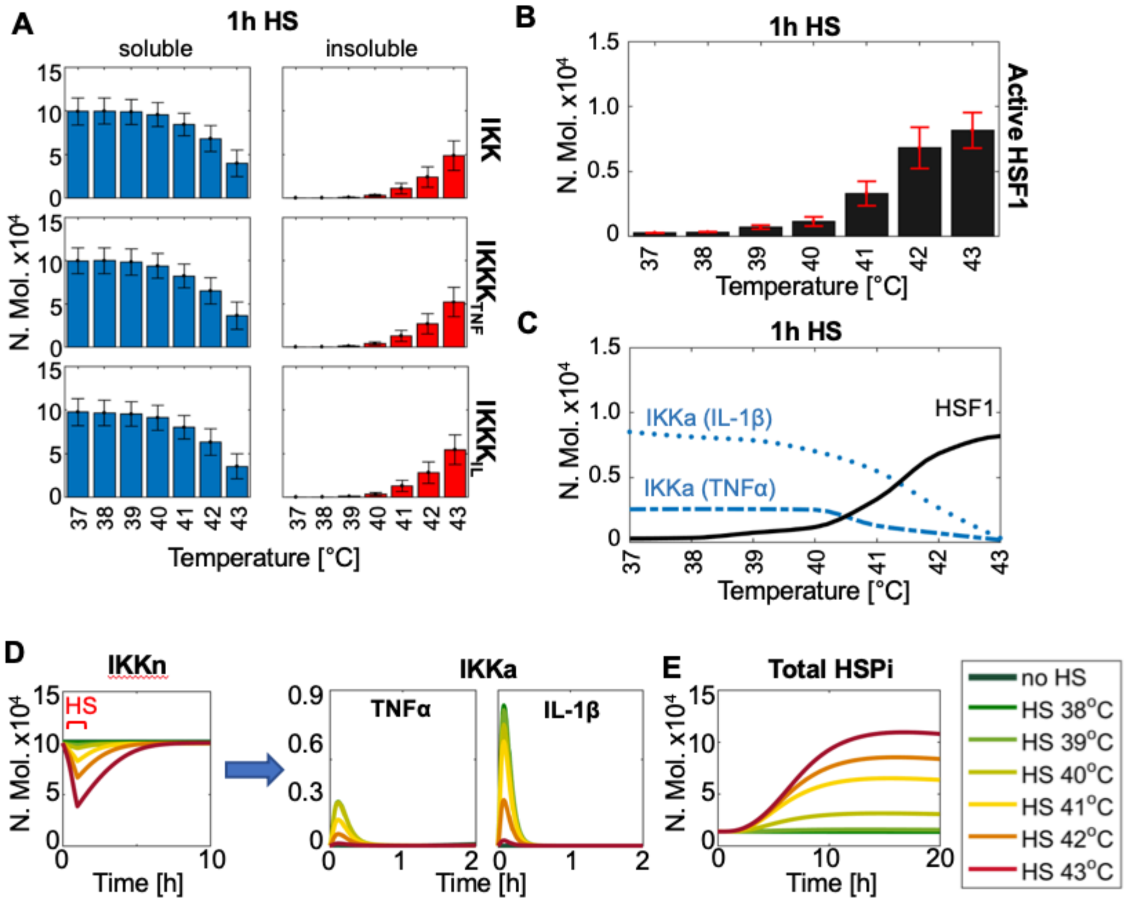


**Figure S5. Temperature sensitivity of the IKK and HSF1 in the mathematical model** (**A**) Comparison of simulated soluble/insoluble IKK and IKKK kinase fractions after 1h HS assuming a 38-43°C temperature range (as indicated on the graph). 37°C represents cells cultured under normal conditions. Shown are average protein levels and standard deviations calculated based on 1,000 single cell model simulations (in number of molecules). (**B**) Simulated level of active HSF1 under conditions as in A. (**C**) A comparison of the peak active IKK kinase level and active HSF1 as a function of temperature. Shown are average protein levels, calculated from 1,000 single cell model simulations (in number of molecules), following TNFα and IL1β treatment immediately after 1h HS exposure. (**D**) Differential cytokine sensitivity to temperature: temperature-dependent depletion of soluble IKK following HS (left) affects TNFα-induced IKK activity (transition from resting inactive, IKKn to active form, IKKa) more than that of IL1β, due to its lower activation amplitude (right). Shown are averages of 1,000 simulated cells (in number of molecules) treated with cytokine immediately after 1h HS exposure to the indicated temperature range. (**E**) Kinetic of HSPi protein accumulation depends on the HS temperature. Shown are average HSPi levels, calculated from 1,000 single cell model simulations after 1h HS at different temperatures.

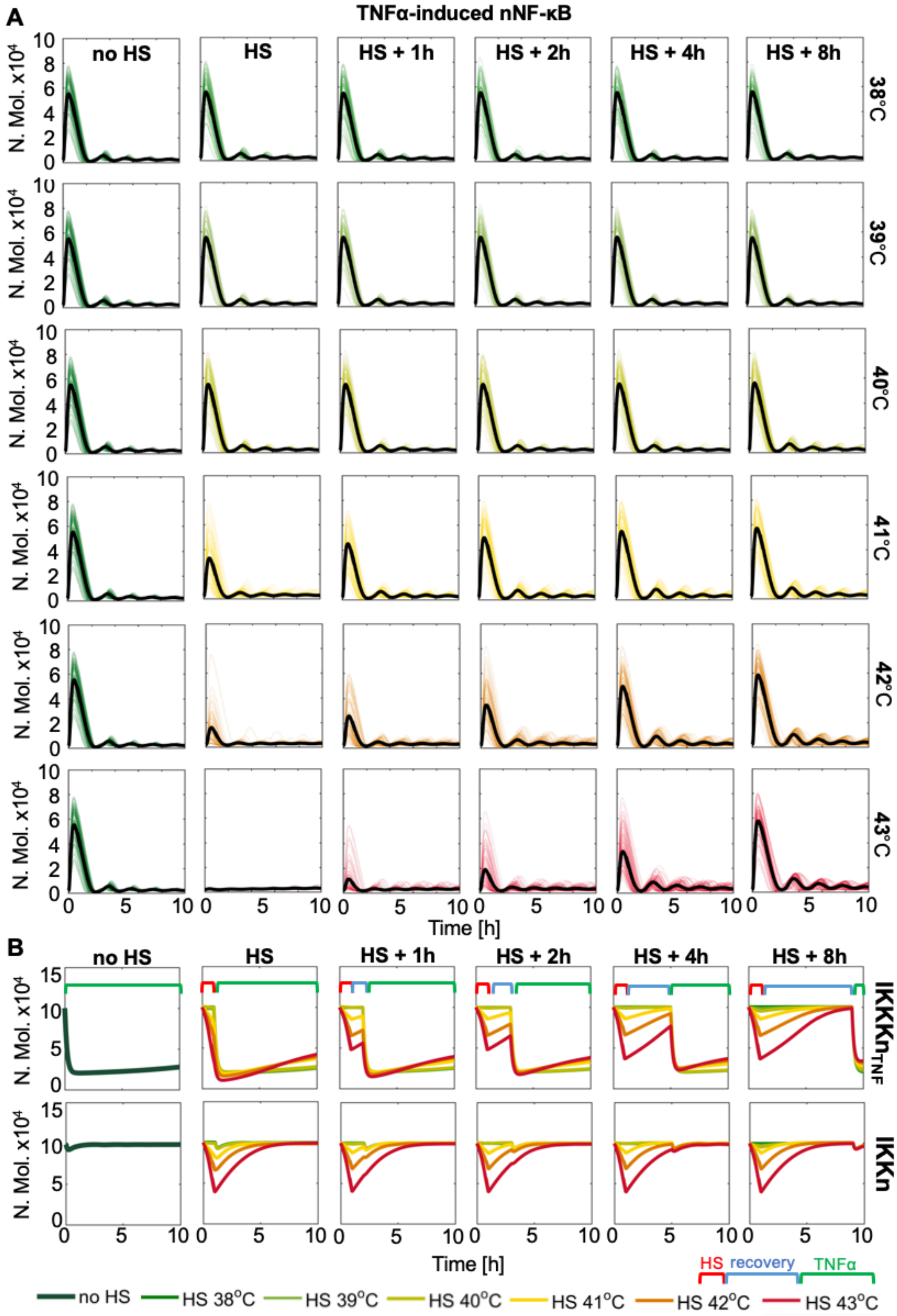


**Figure S6. Model simulations of TNFα-induced responses following range of HS temperatures and different recovery times.** (**A**) Cells are exposed to 1h HS from a temperature range and recovered for up to 8h before cytokine stimulation. Shown are sample 100 time-courses of nuclear NF-κB levels (coloured lines) and average nuclear NF-κB levels (in black), calculated from 1,000 single cell simulations (in number of molecules). (**B**) Comparison of IKK and IKKK kinase levels in simulated data from A.

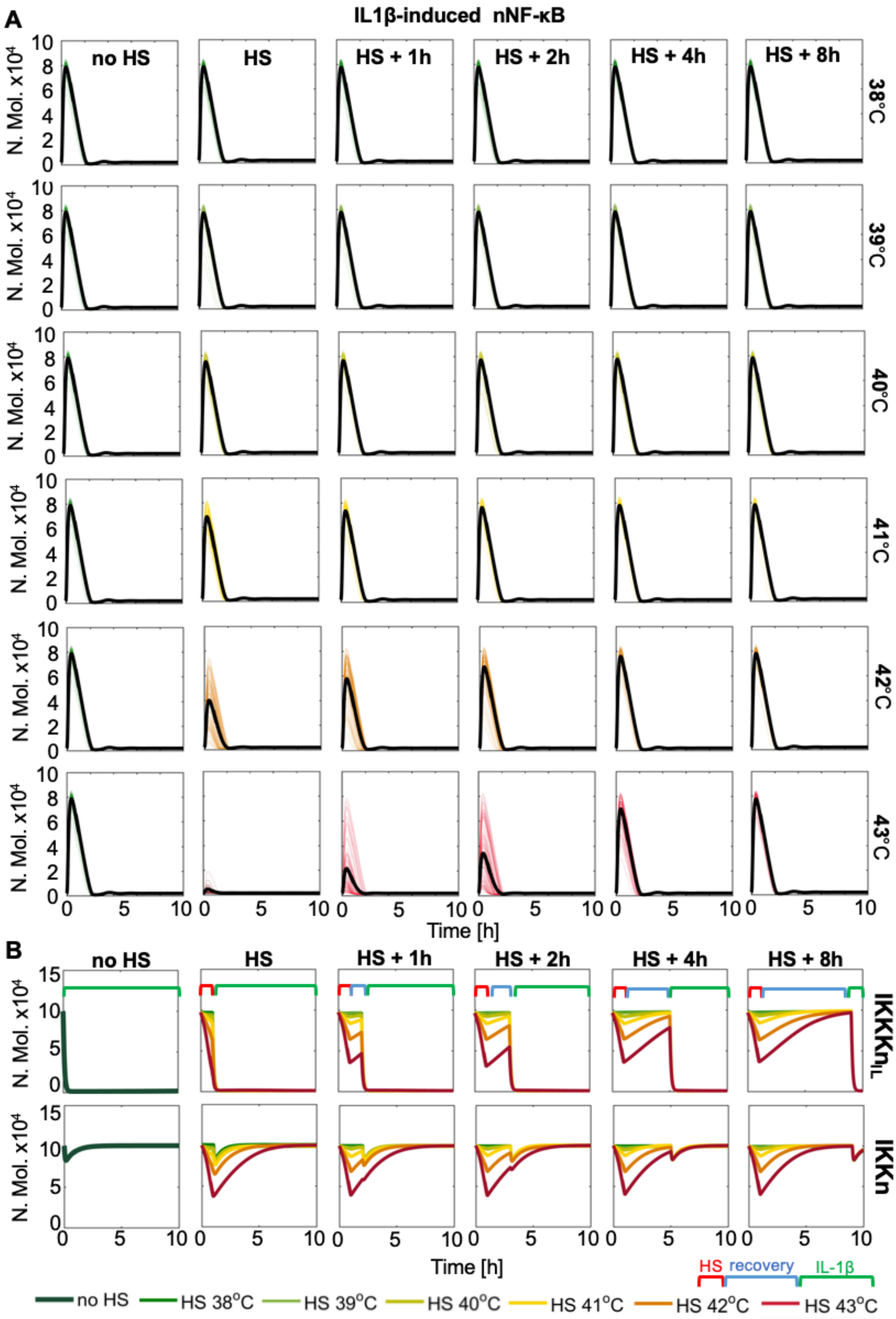


**Figure S7. Model simulations of IL1β-induced responses following range of HS temperatures and different recovery times.** (**A**) Cells are exposed to 1h HS from a temperature range and recovered for up to 8h before cytokine stimulation. Shown are sample 100 time-courses of nuclear NF-κB levels (coloured lines) and average trajectory (in black), calculated from 1,000 single cell simulations (in number of molecules). (**B**) Comparison of IKK and IKKK kinase levels in simulated data from A.

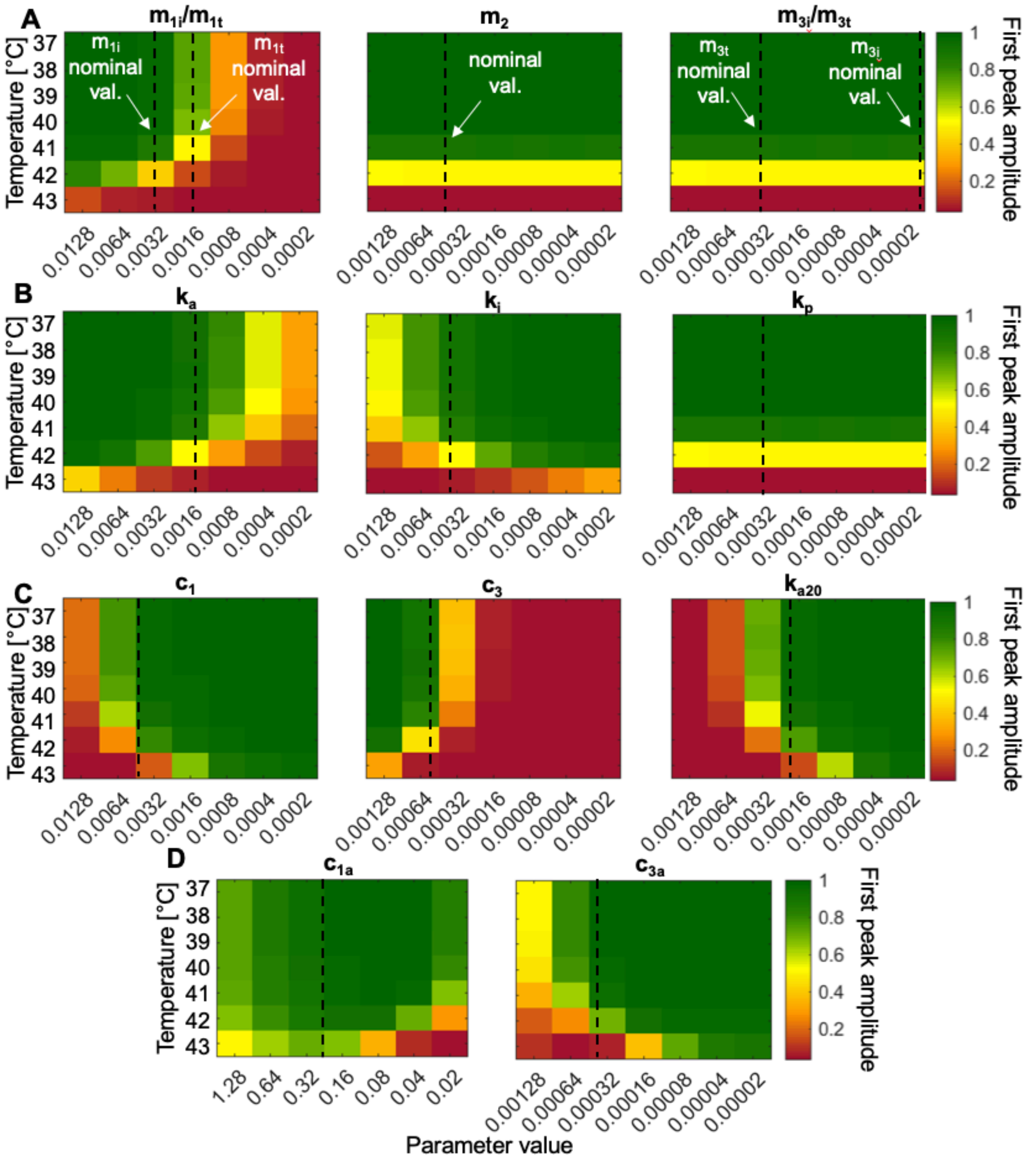


| | | | |
|---|---|---|---|
| A | $m_{1i}/m_{1t}$ | $IKKK_{IL}/IKKK_{TNF}$ activation rate | IKKK module |
| | $m_2$ | IKKK inactivation rate | |
| | $m_{3i}/m_{3t}$ | $IKKK_{IL}/IKKK_{TNF}$ recovery rare | |
| B | $k_a$ | IKK activation rate caused by $IKKK_{IL}/IKKK_{TNF}$ | IKK module |
| | $k_i$ | Spontanouous IKK inactivation | |
| | $k_p$ | Recovery of IKK | |
| C | $c_1$ | Inducible A20 mRNA synthesis | A20 module |
| | $c_3$ | A20 mRNA degradation | |
| | $k_{a20}$ | IKKK degradation rate caused by A20 | |
| D | $c_{1a}$ | Inducible IκB mRNA synthesis | IκB module |
| | $c_{3a}$ | IκB mRNA degradation | |

**Figure S8. Temperature sensitivity analysis of the NF-κB signalling network.** Shown are heat maps describing the influence of model parameters (listed in the table below) involved in (**A**) IKKK, (**B**) IKK, (**C**) A20 and (**D**) IκBα regulation for a range of HS temperatures. All results show sensitivity index calculated for the average nuclear NF-κB levels in the first peak based on 1,000 single cell simulations, normalised to 0-1. Vertical changes indicate increased sensitivity to temperature, nominal parameter values for TNFα and IL1β transduction pathways are indicated with broken lines.

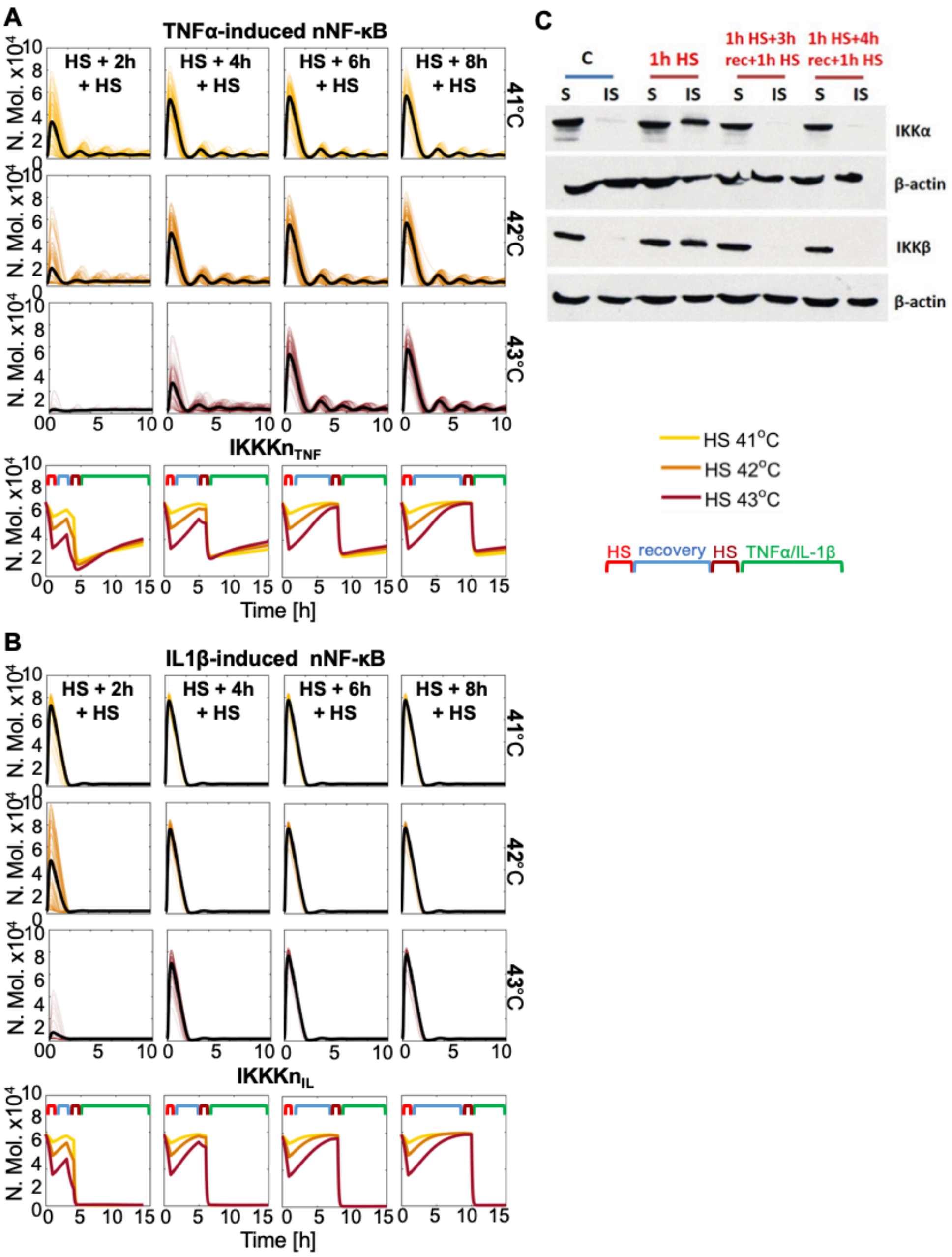


**Figure S9. Responses to repeated HS treatment.** (**A**) Model simulations of cells exposed to repeated 1h HS from a temperature range at a different time interval (from 2 to 8h) and treated with TNFα (immediately after the second HS exposure). Shown are sample 100 time-courses of nuclear NF-κB levels (coloured lines) and average trajectory (in black), calculated from 1,000 single cell simulations across conditions (in number of molecules). Bottom: comparison of the corresponding $IKKK_{TNF}$ kinase levels following different treatment protocols. (**B**) Simulation of responses to IL1β, following the protocol described in A. (**C**) Western blot analysis of soluble (S) and insoluble (IS) IKKα and IKKβ proteins level in MCF7 cells. Cells were either cultured under normal conditions, 37°C, subjected to 1h 43°C HS or subjected to repeated HS after 3 or 4h (as indicated on the graph). β-actin was used as a loading control.

| Gene name | Primer sequence (5’→3’)<br>(L-left, R-right) | Gene ref. number |
|---|---|---|
| *NFKBIA* | L: TGGTGTCCTTGGGTGCTGAT<br>R: GGCAGTCCGGCCATTACA | NM_020529 |
| *TNFAIP3* | L: GCCCTCATCGACAGAAACAT<br>R: CACAAGCTTCCGGACTTCTC | NM_006290 |
| *CCL2* | L: CAAGCAGAAGTGGGTTCAGGAT<br>R: TCTTCGGAGTTTGGGTTTGC | NM_002982.3 |
| *TNFA* | L: CTGCACTTTGGAGTGATCGG<br>R: TCAGCTTGAGGGTTTGCTAC | NM_000594 |
| *HSPA1A* | L: CCGAGAAGGACGAGTTTGAG<br>R: AACAGCAATCTTGGAAAGGC | NM_005345.5 |
| *GAPDH* | L: ACCCAGAAGACTGTGGATGG<br>R: TTCAGCTCAGGGATGACCTT | NM_002046 |

**Table S1**. qRT-PCR primer sequences used in the study.

| | |
|---|---|
| $\mathrm{NF\kappa B}(t)$ | Cytoplasmic amount of free NF-κB |
| $\mathrm{I\kappa B\alpha}(t)$ | Cytoplasmic amount of free IκBα |
| $(\mathrm{I\kappa B\alpha}\vert\mathrm{NF\kappa B})(t)$ | Cytoplasmic amount of NF-κB and IκBα complexes |
| $\mathrm{NF\kappa B}_n(t)$ | Nuclear amount of free NF-κB |
| $\mathrm{I\kappa B\alpha}_n(t)$ | Nuclear amount of free IκBα |
| $(\mathrm{I\kappa B\alpha}_n\vert\mathrm{NF\kappa B}_n)(t)$ | Nuclear amount of NF-κB and IκBα complexes |
| $\mathrm{I\kappa B\alpha}_t(t)$ | Amount of IκBα mRNA transcript |
| $IKKn(t)$ | Cytoplasmic amount of neutral form of IKK kinase |
| $IKKa(t)$ | Cytoplasmic amount of active form of IKK |
| $A20_t(t)$ | Amount of A20 mRNA transcript |
| $A20(t)$ | Cytoplasmic amount of A20 |
| $\mathrm{IKKKn}_{TNF}(t)$ | Cytoplasmic amount of neutral form of $\mathrm{IKKK_{TNF}}$ kinase |
| $\mathrm{IKKKa}_{TNF}(t)$ | Cytoplasmic amount of active form of $\mathrm{IKKK_{TNF}}$ kinase |
| $R_{TNF}(t)$ | Amount of active TNFα receptor |
| $TNF(t)$ | Amount of extracellular TNFα |
| $\mathrm{IKKKn}_{IL}(t)$ | Cytoplasmic amount of neutral form of $\mathrm{IKKK_{IL}}$ kinase |
| $\mathrm{IKKKa}_{IL}(t)$ | Cytoplasmic amount of active form of $\mathrm{IKKK_{IL}}$ kinase |
| $R_{IL}(t)$ | Amount of active IL-1β receptor |
| $IL(t)$ | Amount of extracellular IL-1β |
| $IKK_{IS}(t)$ | Cytoplasmic amount of insoluble IKK kinase |
| $\mathrm{IKKK}_{IL\,IS}(t)$ | Cytoplasmic amount of insoluble $\mathrm{IKKK_{IL}}$ kinase |
| $\mathrm{IKKK}_{TNF\,IS}(t)$ | Cytoplasmic amount of insoluble $\mathrm{IKKK_{TNF}}$ kinase |
| $(HSPc\vert IKK_{IS})(t)$ | Cytoplasmic amount of constitutive HSP and insoluble IKK complexes |
| $(HSPc\vert\mathrm{IKKK}_{IL\,IS})(t)$ | Cytoplasmic amount of constitutive HSP and insoluble $\mathrm{IKKK_{IL}}$ complexes |
| $(HSPi\vert\mathrm{IKKK}_{TNF\,IS})(t)$ | Cytoplasmic amount of inducible HSP and insoluble $\mathrm{IKKK_{TNF}}$ complexes |
| $(HSPi\vert HSF)(t)$ | Amount of inducible HSP and HSF1 complexes |
| $HSF(t)$ | Amount of free HSF1 |
| $HSPi_{tn}(t)$ | Nuclear amount of HSPi mRNA transcript |
| $HSPi_t(t)$ | Cytoplasmic amount of HSPi mRNA transcript |
| $HSPi(t)$ | Amount of free inducible HSP |
| $HSPc(t)$ | Amount of free constitutive HSP |

**Table S2.** Mathematical model variables.

$$\frac{d}{dt}\mathrm{NF\kappa B}(t) = k_{d1a} \times (\mathrm{I\kappa B\alpha|NF\kappa B})(t) - k_{a1a} \times (\mathrm{I\kappa B\alpha|NF\kappa B})(t) - k_{i1} \times \mathrm{NF\kappa B}(t) + k_v \times k_{e1} \times \mathrm{NF\kappa B}_n(t) + k_{t2a} \times (\mathrm{pI\kappa B\alpha|NF\kappa B})(t) \quad (1)$$

$$\frac{d}{dt}\mathrm{I\kappa B\alpha}(t) = k_{d1a} \times (\mathrm{I\kappa B\alpha|NF\kappa B})(t) - k_{a1a} \times (\mathrm{I\kappa B\alpha|NF\kappa B})(t) - k_{i3a} \times \mathrm{I\kappa B\alpha}(t) + k_v \times k_{e3a} \times \mathrm{I\kappa B\alpha}_n(t) - c_{4a} \times \mathrm{I\kappa B\alpha}(t) + c_{2a} \times \mathrm{I\kappa B\alpha}_t(t) - k_{c1a} \times IKKa(t) \times \mathrm{I\kappa B\alpha}(t) \quad (2)$$

$$\frac{d}{dt}(\mathrm{I\kappa B\alpha|NF\kappa B})(t) = k_{a1a} \times (\mathrm{I\kappa B\alpha|NF\kappa B})(t) - k_{d1a} \times (\mathrm{I\kappa B\alpha|NF\kappa B})(t) + k_v \times k_{e2a} \times (\mathrm{I\kappa B\alpha}_n|\mathrm{NF\kappa B}_n)(t) - k_{c2a} \times IKKa(t) \times (\mathrm{I\kappa B\alpha|NF\kappa B})(t) \quad (3)$$

$$\frac{d}{dt}\mathrm{NF\kappa B}_n(t) = k_{d1a} \times (\mathrm{I\kappa B\alpha}_n|\mathrm{NF\kappa B}_n) - k_{a1a} \times k_v \times \mathrm{I\kappa B\alpha}_n(\mathrm{t}) \times \mathrm{NF\kappa B}_n(t) + k_{i1} \times \mathrm{NF\kappa B}(t) - k_v \times k_{e1} \times \mathrm{NF\kappa B}_n(t) \quad (4)$$

$$\frac{d}{dt}\mathrm{I\kappa B\alpha}_n(t) = k_{d1a} \times (\mathrm{I\kappa B\alpha}_n|\mathrm{NF\kappa B}_n)(t) - k_v \times k_{a1a} \times \mathrm{I\kappa B\alpha}_n(\mathrm{t}) \times \mathrm{NF\kappa B}_n(t) + k_{i3a} \times \mathrm{I\kappa B\alpha}(t) - k_v \times k_{e3a} \times \mathrm{I\kappa B\alpha}_n(t) - c_{4a} \times \mathrm{I\kappa B\alpha}_n(t) \quad (5)$$

$$\frac{d}{dt}(\mathrm{I\kappa B\alpha}_n|\mathrm{NF\kappa B}_n)(t) = k_v \times k_{a1a} \times \mathrm{I\kappa B\alpha}_n(\mathrm{t}) \times \mathrm{NF\kappa B}_n(t) - k_{d1a} \times (\mathrm{I\kappa B\alpha}_n|\mathrm{NF\kappa B}_n)(t) - k_v \times k_{e3a} \times (\mathrm{I\kappa B\alpha}_n|\mathrm{NF\kappa B}_n)(t) \quad (6)$$

$$\frac{d}{dt}\mathrm{I\kappa B\alpha}_t(t) = c_{1a}\frac{\mathrm{NF\kappa B}_n(t)^h}{\mathrm{NF\kappa B}_n(t)^h + k^h} - c_{3a} \times \mathrm{I\kappa B\alpha}_t(t) \quad (7)$$

$$\frac{d}{dt}IKKn(t) = k_p(1 + K_r \times HSPc(\mathrm{t})) \times \left(IKKtot - IKKn(\mathrm{t}) - IKKi(\mathrm{t}) - IKK_{IS}(\mathrm{t}) - (HSPc|IKK_{IS})(\mathrm{t})\right) - k_a \frac{\mathrm{IKKKa}_{TNF}(t)^{ha}}{\mathrm{IKKKa}_{TNF}(t)^{ha} + sIKKK^{ha}} IKKn(\mathrm{t}) - k_a \frac{\mathrm{IKKKa}_{IL}(t)^{ha}}{\mathrm{IKKKa}_{IL}(t)^{ha} + sIKKK^{ha}} IKKn(\mathrm{t}) - (Temp - 37)^n \times \frac{k_{IS1}}{1+\left(\frac{HSPi(t)+HSPc(t)}{Kh}\right)^{n_2}} \times IKKn(t) + k_{re1} \times (HSPc|IKK_{IS})(\mathrm{t}) \quad (8)$$

$$\frac{d}{dt}IKKa(t) = k_a \frac{\mathrm{IKKKa}_{TNF}(t)^{ha}}{\mathrm{IKKKa}_{TNF}(t)^{ha} + sIKKK^{ha}} IKKn(\mathrm{t}) + k_a \frac{\mathrm{IKKKa}_{IL}(t)^{ha}}{\mathrm{IKKKa}_{IL}(t)^{ha} + sIKKK^{ha}} IKKn(\mathrm{t}) - k_i \times IKKa(t) - (Temp - 37)^n \times \frac{k_{IS1}}{1+\left(\frac{HSPi(t)+HSPc(t)}{Kh}\right)^{n_2}} \times IKKa(t) \quad (9)$$

$$\frac{d}{dt}A20_t(t) = A_{A20} \times c_1 \frac{\mathrm{NF\kappa B}_n(t)^h}{\mathrm{NF\kappa B}_n(t)^h + k^h} - c_3 \times A20_t(t) \quad (10)$$

$$\frac{d}{dt}A20(t) = A_{A20} \times c_2 \times A20_t(t) - c_4 \times A20(t) \quad (11)$$

$$\frac{d}{dt}\mathrm{IKKKn}_{TNF}(t) = m_{3t}(1 + K_r \times HSPi(\mathrm{t})) \times \left(IKKKtot_{TNF} - \mathrm{IKKKn}_{TNF}(t) - \mathrm{IKKKa}_{TNF}(t) - \mathrm{IKKK}_{TNF\,IS}(t) - (HSPi|\mathrm{IKKK}_{TNF\,IS})(t)\right) - m_{1t}\frac{R_{TNF}(t)}{R_{TNF}(t)+s_t}\mathrm{IKKKn}_{TNF}(t) - (Temp - 37)^n \times \frac{k_{IS3}}{1+\left(\frac{HSPi(t)+HSPc(t)}{Kh}\right)^{n_2}} \times \mathrm{IKKKn}_{TNF}(t) + k_{re3} \times (HSPi|\mathrm{IKKK}_{TNF\,IS})(\mathrm{t}) \quad (12)$$

$$\frac{d}{dt}\mathrm{IKKKa}_{TNF}(t) = m_{1t}\frac{R_{TNF}(t)}{R_{TNF}(t)+s_t}\mathrm{IKKKn}_{TNF}(t) - \left(m_2 + k_{a20} \times A20(t)\right) \times \mathrm{IKKKa}_{TNF}(t) - (Temp - 37)^n \times \frac{k_{IS3}}{1+\left(\frac{HHSPi(t)+HSPc(t)}{Kh}\right)^{n_2}} \times \mathrm{IKKKa}_{TNF}(t) \quad (13)$$

$$\frac{d}{dt}R_{TNF}(t) = r_1 \times TNF(t) \times \left(Rtot - R_{TNF}(t)\right) - r_2 \times R_{TNF}(t) \quad (14)$$

$$\frac{d}{dt}TNF(t) = -c_5 \times TNF(t) - r_1 \times TNF(t) \times \left(Rtot - R_{TNF}(t)\right) \tag{15}$$

$$\frac{d}{dt}\mathrm{IKKKn}_{IL}(t) = m_{3i}(1 + K_r \times HSPc(\mathrm{t})) \times \left(IKKKtot_{IL} - \mathrm{IKKKn}_{IL}(t) - \mathrm{IKKKa}_{IL}(t) - \mathrm{IKKK}_{IL\ IS}(t) - (HSPc|\mathrm{IKKK}_{IL\ IS})(t)\right) - m_{1i}\frac{R_{IL}(t)}{R_{IL}(t)+s_t}\mathrm{IKKKn}_{IL}(t) - (Temp - 37)^n \times \frac{k_{IS2}}{1+\left(\frac{HSPi(t)+HSPc(t)}{Kh}\right)^{n_2}} \times \mathrm{IKKKn}_{IL}(t) + k_{re2} \times (HSPc|\mathrm{IKKK}_{IL\ IS})(\mathrm{t}) \tag{16}$$

$$\frac{d}{dt}\mathrm{IKKKa}_{IL}(t) = m_{1i}\frac{R_{IL}(t)}{R_{IL}(t)+s_t}\mathrm{IKKKn}_{IL}(t) - \left(m_2 + k_{a20} \times A20(t)\right) \times \mathrm{IKKKa}_{IL}(t) - (Temp - 37)^n \times \frac{k_{IS2}}{1+\left(\frac{HSPi(t)+HSPc(t)}{Kh}\right)^{n_2}} \times \mathrm{IKKKa}_{IL}(t) \tag{17}$$

$$\frac{d}{dt}R_{IL}(t) = r_1 \times IL(t) \times \left(Rtot - R_{IL}(t)\right) - r_2 \times R_{IL}(t) \tag{18}$$

$$\frac{d}{dt}IL(t) = -c_5 \times IL(t) - r_1 \times IL(t) \times \left(Rtot - R_{IL}(t)\right) \tag{19}$$

$$\frac{d}{dt}IKK_{IS}(t) = (Temp - 37)^n \times k_{IS1} \times IKKn(t) + (Temp - 37)^n \times k_{IS1} \times IKKa(t) - k_{hs1}\frac{HSPc(t)}{K_m+HSPc(t)}IKK_{IS}(t) \tag{20}$$

$$\frac{d}{dt}\mathrm{IKKK}_{TNF\ IS}(t) = (Temp - 37)^n \times k_{IS3} \times \mathrm{IKKKn}_{TNF}(t) + (Temp - 37)^n \times k_{IS3} \times \mathrm{IKKKa}_{TNF}(t) - k_{hs3}\frac{HSPi(t)}{K_m+HSPi(t)}\mathrm{IKKK}_{TNF\ IS}(t) \tag{21}$$

$$\frac{d}{dt}\mathrm{IKKK}_{IL\ IS}(t) = (Temp - 37)^n \times k_{IS2} \times \mathrm{IKKKn}_{IL}(t) + (Temp - 37)^n \times k_{IS2} \times \mathrm{IKKKa}_{IL}(t) - k_{hs2}\frac{HSPc(t)}{K_m+HSPc(t)}\mathrm{IKKK}_{IL\ IS}(t) \tag{22}$$

$$\frac{d}{dt}(HSPc|IKK_{IS})(t) = k_{hs1}\frac{HSPc(t)}{K_m+HSPc(t)}IKK_{IS}(t) - k_{re1} \times (HSPc|IKK_{IS})(\mathrm{t}) \tag{23}$$

$$\frac{d}{dt}(HSPi|\mathrm{IKKK}_{TNF\ IS})(t) = k_{hs3}\frac{HSPi(t)}{K_m+HSPi(t)}\mathrm{IKKK}_{TNF\ IS}(t) - k_{re3} \times (HSPi|\mathrm{IKKK}_{TNF\ IS})(\mathrm{t}) \tag{24}$$

$$\frac{d}{dt}(HSPc|\mathrm{IKKK}_{IL\ IS})(t) = k_{hs2}\frac{HSPc(t)}{K_m+HSPc(t)}\mathrm{IKKK}_{IL\ IS}(t) - k_{re2} \times (HSPc|\mathrm{IKKK}_{IL\ IS})(\mathrm{t}) \tag{25}$$

$$\frac{d}{dt}(HSPi|HSF)(t) = -k_{a1} \times (HSPi|HSF)(t) - k_{a2} \times (HSPi|HSF)(t) \times \mathrm{IKKK}_{TNF\ IS}(t) + k_{ia}\frac{HSPi(t)}{K_m+HSPi(t)}HSF(t) \tag{26}$$

$$\frac{d}{dt}HSF(t) = k_{a1} \times (HSPi|HSF)(t) + k_{a2} \times (HSPi|HSF)(t) \times \mathrm{IKKK}_{TNF\ IS}(t) - k_{ia}\frac{HSPi(t)}{K_m+HSPi(t)}HSF(t) \tag{27}$$

$$\frac{d}{dt}HSPi_{tn}(t) = k_{p1}\frac{HSF(t)^{hi}}{HSF(t)^{hi}+k^{hi}} - k_t \times HSPi_t \tag{28}$$

$$\frac{d}{dt}HSPi_t(t) = k_t \times HSPi_t(t) - k_{d1} \times HSPi_{tc}(t) \tag{29}$$

$$\frac{d}{dt}HSPi(t) = k_{p2} \times HSPi_{tc}(t) - k_{d2} \times HSPi(t) - k_{ia}\frac{HSPi(t)}{K_m + HSPi(t)}HSF(t) - k_{hs3}\frac{HSPi(t)}{K_m + HSPi(t)}\mathrm{IKKK}_{TNF\ IS}(t) + k_{a1} \times (HSPi|HSF)(t) + k_{re3} \times (HSPi|\mathrm{IKKK}_{TNF\ IS})(\mathrm{t}) \quad (30)$$

$$\frac{d}{dt}HSPc(t) = -k_{hs1}\frac{HSPc(t)}{K_m + HSPc(t)}IKK_{IS}(t) + k_{re1} \times (HSPc|IKK_{IS})(\mathrm{t}) - k_{hs2}\frac{HSPc(t)}{K_m + HSPc(t)}\mathrm{IKKK}_{IL\ IS}(t) + k_{re2} \times (HSPc|\mathrm{IKKK}_{IL\ IS})(\mathrm{t}) \quad (31)$$

**Table S3.** Mathematical model equations.

| Symbol | Value | Reference | Symbol | Value | Reference |
|---|---|---|---|---|---|
| $k_v$ | 3.3 | [1] | $m_{3t}$ | $2.94\cdot10^{-4}$ [$s^{-1}$] | Fitted |
| $c_1$ | 0.005 [$s^{-1}$] | [1] | $r_1$ | $4\cdot10^{-9}$ [$s^{-1}$] | [1] |
| $c_{1a}$ | 0.0286 [$s^{-1}$] | Fitted | $R_2$ | 0.0032 [$s^{-1}$] | [1] |
| h | 2 | [1] | Rtot | 2500 | [1] |
| $c_2$ | 0.5 [$s^{-1}$] | [1] | st | 50 | [1] |
| $c_{2a}$ | 0.5 [$s^{-1}$] | [1] | $k_{e1}$ | $5.20\cdot10^{-5}$ [$s^{-1}$] | [1] |
| $c_5$ | 0.000037 [$s^{-1}$] | [1] | $k_{e3a}$ | $5.00\cdot10^{-4}$ [$s^{-1}$] | [1] |
| $k_{a1a}$ | $8.0\cdot10^{-7}$ [$s^{-1}$] | [1] | $k_{IS1}$ | $9.0\cdot10^{-8}$ [$s^{-1}$] | Fitted |
| $k_{d1a}$ | $8.0\cdot10^{-4}$ [$s^{-1}$] | [1] | $k_{IS2}$ | $9.0\cdot10^{-8}$ [$s^{-1}$] | Assumed |
| $c_{3a}$ | $4.4\cdot10^{-4}$ | [1] | $k_{IS3}$ | $9.0\cdot10^{-8}$ [$s^{-1}$] | Assumed |
| $c_{4a}$ | $7.6\cdot10^{-4}$ | [1] | $k_{hs1}$ | $6.9\cdot10^{-4}$ [$s^{-1}$] | Fitted |
| $c_3$ | $6.6\cdot10^{-4}$ | [1] | $k_{hs2}$ | $4.9\cdot10^{-4}$ [$s^{-1}$] | Assumed |
| $c_4$ | $7.5\cdot10^{-4}$ | [1] | $k_{hs3}$ | $1.0\cdot10^{-4}$ [$s^{-1}$] | Assumed |
| $k_{i1}$ | 0.0026 [$s^{-1}$] | [1] | $k_{re1}$ | $4.9\cdot10^{-4}$ [$s^{-1}$] | Fitted |
| $k_{i3a}$ | 0.001 [$s^{-1}$] | [1] | $k_{re2}$ | $3.5\cdot10^{-4}$ [$s^{-1}$] | Assumed |
| $k_{e2a}$ | 0.01 [$s^{-1}$] | [1] | $k_{re3}$ | $2.3\cdot10^{-4}$ [$s^{-1}$] | Assumed |
| k | $2.111\cdot10^{3}$ | [1] | Km | 10000 | Assumed |
| $k_{a20}$ | 0.00023 [$s^{-1}$] | [1] | n | 3 | Fitted |
| $k_{c1a}$ | $9.8234\cdot10^{-8}$ [$s^{-1}$] | [1] | $k_{a1}$ | $0.7\cdot10^{-5}$ [$s^{-1}$] | Assumed |
| $k_{c2a}$ | $8.5510\cdot10^{-7}$ [$s^{-1}$] | [1] | $k_{a2}$ | $3.5\cdot10^{-8}$ [$s^{-1}$] | Fitted |
| $k_{t2a}$ | 0.1 [$s^{-1}$] | [1] | $k_{ia}$ | $9.1\cdot10^{-4}$ [$s^{-1}$] | Fitted |
| $k_p$ | 0.00033 [$s^{-1}$] | [1] | $k_{p1}$ | 0.0026 [$s^{-1}$] | [3] |
| $k_a$ | 0.0014 [$s^{-1}$] | [1] | $k_t$ | $0.7\cdot10^{-4}$ [$s^{-1}$] | Assumed |
| $k_i$ | 0.0040 [$s^{-1}$] | [1] | $k_{d1}$ | $2.16\cdot10^{-4}$ [$s^{-1}$] | [3] |
| ha | 3 | [1] | $k_{p2}$ | 0.50 [$s^{-1}$] | Fitted |
| $m_{1i}$ | 0.0033 [$s^{-1}$] | Fitted | $k_{d2}$ | $4.36\cdot10^{-6}$ [$s^{-1}$] | [2] |
| $m_{1t}$ | 0.0015 [$s^{-1}$] | Fitted | hi | 3 | [4] |
| sIKKK | 350 | [1] | $K_r$ | $1.43\cdot10^{-5}$ [$s^{-1}$] | Assumed |
| $m_2$ | 0.0005 [$s^{-1}$] | [1] | $n_2$ | 3 | Assumed |
| $m_{3i}$ | $1.43\cdot10^{-5}$ [$s^{-1}$] | Fitted | Kh | $1.63\cdot10^{4}$ | Assumed |

**Table S4.** Model parameters.
